# ASPARAGINE-RICH METASTATIC NICHES DRIVE PROSTATE CANCER ORGANOTROPISM BY ENABLING TRANSLATIONAL REWIRING TOWARD N-GLYCOSYLATED PROTEINS

**DOI:** 10.64898/2026.02.27.708521

**Authors:** E. Pranzini, L. Ippolito, M. Iozzo, S. Romagnoli, G. Bertoli, G. Venditti, M. Lulli, A. Santi, G. Comito, S. Polvani, T. Lottini, M. Benelli, D. Scumaci, P. Chiarugi, E. Giannoni

## Abstract

Metastatic disease is the leading cause of mortality in prostate cancer (PC), with bone as the preferential site of dissemination and the lung frequently affected secondarily. Metabolic interactions between disseminated tumor cells and tissue-specific microenvironments play a key role in shaping site-specific metastatic patterns. In particular, the availability of certain metabolites before, during, and after the establishment of overt metastases represents a critical determinant of colonization. Here we show that asparagine (Asn) is selectively enriched in bone and lung microenvironments and supports PC metastatic colonization. Dietary Asn restriction selectively impairs bone and lung metastases *in vivo*, without affecting metastatic burden in other organs. Mechanistically, disseminated PC cells arriving at Asn-rich niches rely on extracellular Asn due to decreased asparagine synthetase (ASNS) expression enabling the activation of mTORC1-dependent translational program. Elevated Asn availability selectively promotes the synthesis of Asn-rich proteins enriched in N-glycosylation motifs (Asn-X-Ser/Thr), leading to enhanced global protein N-glycosylation during early metastatic colonization. This metabolic adaptation facilitates cell-cell interactions and promotes adhesion to bone- and lung-specific extracellular matrices. Among the N-glycosylated proteins induced by Asn, CD44 emerges as a central effector of PC bone metastases. Extracellular Asn shortening, genetic or pharmacological disruption N-glycosylation, or silencing of CD44 abolish the pro-metastatic advantage conferred by Asn. Together, these findings identify environmental Asn as a niche-specific metabolic cue that drives PC organotropism by rewiring translation toward proteins enriched in N-glycosylation sites, thereby enhancing adhesive interactions and revealing metabolic vulnerabilities that could be therapeutically exploited to interfere with earliest steps of metastatic colonization in PC.

## INTRODUCTION

While localized prostate cancer (PC) is often successfully managed, metastases develop in approximately 30% of patients and are the leading cause of death. Bone is the primary site of PC metastasis, with an incidence of nearly 90% of patients. Bone metastasis has been found to aid cancer cell dissemination to other organs during metastatic PC evolution, as observed by post-mortem tissue examination mostly indicating lung as secondary preferential metastatic site (40-45% incidence)^1^ over the liver (25%) and brain (9%) ^2^. The determinants of organ-specific metastatic distribution are multifactorial and include vascular accessibility, chemokine signaling, stromal support, and extracellular matrix (ECM) composition^3^. More recently, the metabolic landscape of target organs has emerged as a critical regulator of metastatic seeding and outgrowth^4^. Nutrient availability differs across organ niches, forcing disseminated tumor cells (DTCs) to adopt context-specific metabolic strategies to exploit locally available substrates. Throughout the metastatic cascade, DTCs experience profound metabolic remodeling to overcome stress. Upon arrival at distant sites, they must further adjust to the new environment to enable successful outgrowth. On one side, nutrient scarcity at distant sites imposes selective pressures that necessitate metabolic adaptations in metastatic cells. For example, cancer cells metastasizing the brain compensate local serine and glycine limiting availability by upregulating *de novo* serine synthesis pathway to meet increased demand for nucleotides, ultimately affecting cancer cell proliferation^5^.

Conversely, the abundance of specific metabolites in the metastatic niche can create adaptive opportunities for DTCs for succeeding in metastatic seeding and outgrowth. For example, metastatic breast cancer cells exploit the high levels of pyruvate present in the lung microenvironment^6^ to enhance anaplerosis^7^, activate the mammalian target of rapamycin complex 1 (mTORC1) signaling^8^, and increase α-ketoglutarate production, which in turn activates collagen hydroxylation and maturation^9^, thereby promoting successful metastatic outgrowth. Specifically, increasing evidence demonstrates that the enrichment of specific amino acids within permissive niches can actively promote early metastatic colonization by enabling metabolic adaptations and facilitating metastatic seeding^10^. Through a distinct adaptive strategy, extracellular aspartate (Asp) in the lung activates NMDA receptor–dependent signaling in metastatic breast cancer cells, thereby triggering a translational program that promotes TGFβ-driven collagen synthesis and optimal extracellular matrix remodeling^11^.

In this study, we explored the hypothesis that PC cells colonizing distant tissues, while adapting to niche-specific metabolic challenges (such as hypoxia^12^ and limited nutrient availability in the bone microenvironment^13^) also exploit locally enriched metabolites to support metastatic establishment. Comparative analysis of the metabolic landscapes of organs most frequently targeted by PC metastases allowed us to identify asparagine (Asn) as a key metabolic determinant specifically enriched in bone and lung underlying the preferential distribution of metastatic PC cells. Specifically, we demonstrated that high Asn availability within these metastatic niches supports the increased demand for synthesis of proteins enriched in N-glycosylation motifs (Asn-X-Ser/Thr), which are critical for re-establishing cell-cell and cell-ECM interactions during the preliminary stages of tissue colonization. This Asn-driven metabolic adaptation enhances PC cell adhesion to bone and lung matrices, thereby facilitating metastatic seeding and subsequent outgrowth. Notably, we identified CD44 as a central mediator of Asn-mediated adaptations in metastatic cells, highlighting its potential as a therapeutic target to interfere with PC metastatic colonization.

## RESULTS

### 1. Tissues metabolic profiling identifies Asn enrichment in bone and lung as a driver of metastatic organotropism in PC

To profile the metabolic composition of tissues that, to different extents, are recognized as PC metastatic sites, we performed untargeted LC-MS analysis on bone, lung, liver and brain collected from healthy mice. Principal component analysis (PCA) of the global metabolomic data revealed distinct tissue-specific clustering, with bone and lung metabolomes showing a high degree of similarity compared with those of the other tissues (Fig. 1A). Consistently, untargeted LC-MS analysis of tissue extracellular environments (bone marrow and lung interstitial fluid), together with plasma, revealed high correlations across biological replicates, further confirming a clear metabolic resemblance between bone and lung sites. (Suppl. Fig.1A). This pattern suggested that the preferential and sequential colonization of bone and lung by metastatic PC cells might be supported by the metabolic similarities shared by these tissues^14^. Among the metabolic pathways implicated in metastatic progression, amino acid metabolism is increasingly recognized as a key contributor to disseminated tumor cell fitness, supporting their colonization and survival within the metastatic niche^5,10,11^. We therefore focused on investigating the contribution of amino acids to PC metastasis. Using GC-MS analysis, we compared amino acid abundance across multiple tissues, using the prostate as a reference. Notably, Asn emerged as the most consistently enriched amino acid in both bone and lung, whereas its levels were lower in other organs, including brain and liver (Fig. 1B, Suppl. Fig. 1B).

**Fig. 1.**
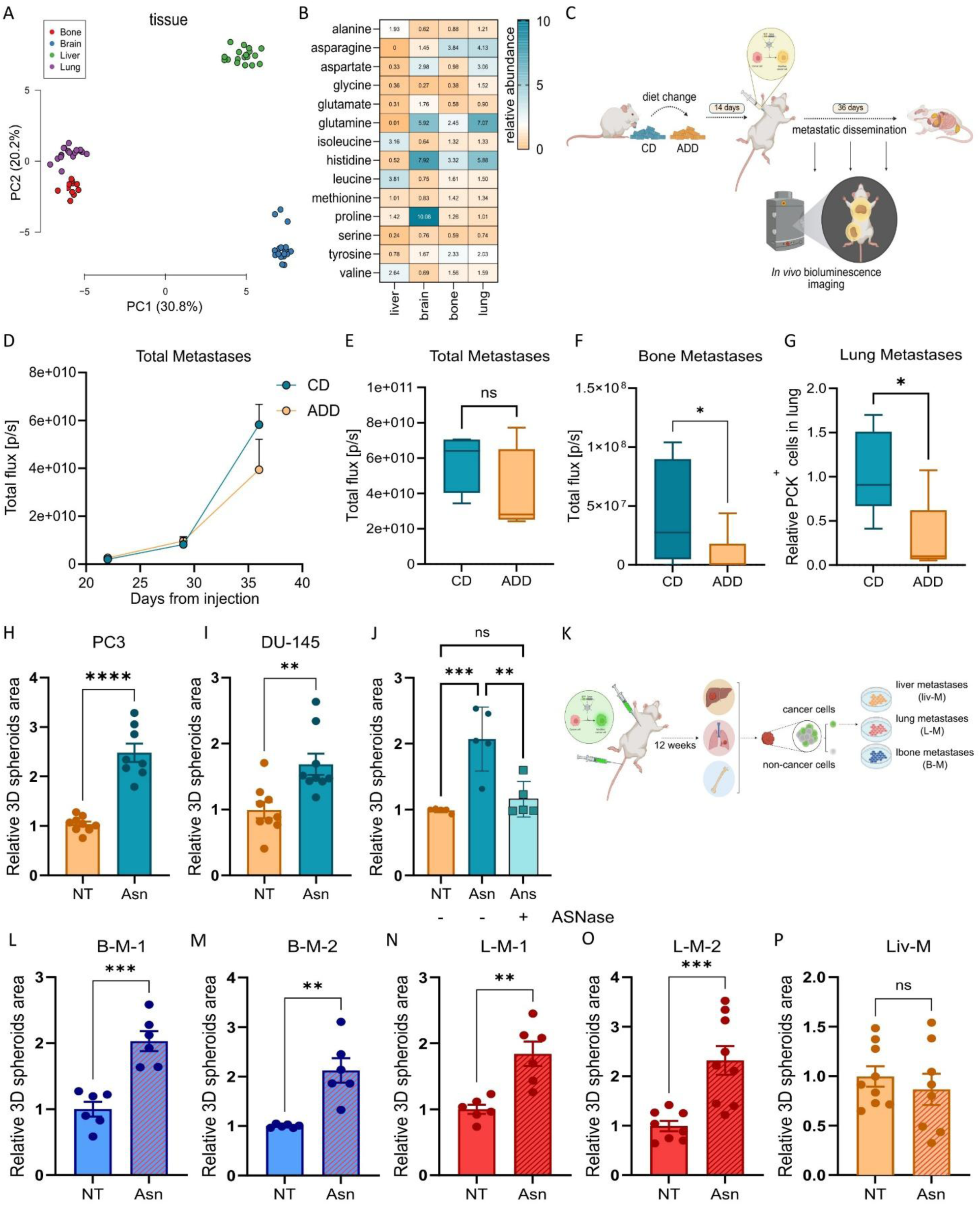
High exogenous Asn enhances the metastatic potential of prostate cancer (PC) cells by sustaining colonization of bone and lung tissues. **(A)** Principal component analysis (PCA) of global metabolomic profiles from tissues (bone, brain, liver, lung) collected from healthy NOD-SCID mice (n = 6) and analyzed by LC-MS. The PCA scatter plot shows PC1 (x-axis), accounting for 30.8% of the total variance, and PC2 (y-axis), explaining 20.2% of the variance. **(B)** Relative amino acid content in tissues from healthy NOD-SCID mice (n=3), measured by GC-MS and normalized to the corresponding amino acid levels in prostate tissue. Each heatmap cell is labeled with the average value. **(C)** Schematic overview of the *in vivo* experimental design used in panels E-G. NOD-SCID mice were conditioned for two weeks with either a normal diet (ND) or an Asn-deprived diet (ADD), followed by intracardiac (i.c.) injection of luciferase-expressing PC3 cells (Luc⁺ PC3). Metastatic progression was monitored longitudinally using the IVIS Lumina S5 system, starting at day 29 post-injection. **(D)** Longitudinal monitoring of metastatic progression using the IVIS Lumina S5 system beginning at day 29 post-injection. Metastatic burden is expressed as total bioluminescent signal (photons/s) per mouse (n = 4). **(E–F)** Quantification of total (E) and bone (F) metastases at day 36 following cancer cell injection, expressed as bioluminescent signal (photons/s). Bone metastases were quantified by measuring signal within bone-specific ROIs as indicated in Suppl. Fig.1E. Welch’s t-test. (n = 4) **(G)** Quantification of lung metastases at the experimental endpoint, expressed as the number of pancytokeratin (PCK)-positive cells per lung section. Welch’s t-test. (n = 4) **(H–I)** Total spheroid area of PC3 (H) and DU145 (I) 3D-C grown with or without Asn (0.1 mM). Values are normalized to control. Unpaired Student’s t-test. **(J)** Total spheroid area of PC3 3D-C cultured with or without Asn (0.1 mM) in the presence or absence of L-asparaginase (ASNase, 0.25 U/ml). One-way ANOVA with Sidak’s correction. **(K)** Schematic overview of *in vivo* selection of metastatic subpopulations isolated from bone, lung, and liver of NOD-SCID mice previously injected i.c. or via the caudal artery (c.a.) with GFP⁺ PC3 cells. Twelve weeks post-injection, tissues were dissociated and GFP⁺ cells were isolated by FACS. **(L–P)** Total spheroid area of bone-derived (L, M), and lung-derived (N, O) liver-derived (P) metastatic cells (obtained after c.a. or i.c. injection, respectively) grown in 3D-C cultures with or without Asn (0.1 mM). Values are normalized to control. Unpaired Student’s t-test. ns, not significant; *P < 0.05; **P < 0.01; ***P < 0.001. Data represent mean ± s.e.m. from at least three independent experiments.

To understand whether Asn availability is really beneficial for the metastatic colonization *in vivo*, mice were fed either with an Asn-deprived diet (ADD) or a control diet (CD) starting from two weeks prior to intracardiac injection (i.c.) of luciferase-expressing PC cells (Luc⁺PC3) allowing for a multi-organ metastasis dissemination. Metastatic progression was monitored from day 29 post-injection (Fig. 1C). No significant differences in body weight or food intake were observed between the two groups (Suppl. Fig. 1C, D). Strikingly, mice maintained on the ADD diet exhibited a slight reduction in the total metastatic burden compared with CD-fed animals (Fig. 1D-E; Suppl. Fig. 1E). However, analysis of regional luminescence signals revealed a selective and significant decrease in bone metastases in the ADD group (Fig. 1F), paralleled by a similar reduction in the total amount of metastatic lesions in the lung (Fig. 1G; Suppl. Fig. 1F). Interestingly, in our model, the ADD regimen did not significantly alter the burden of abdominal metastases compared to CD (Suppl. Fig. 1g). These results suggest that Asn availability dictates PC metastatic tropism and outgrowth specifically in bone and lung.

We first asked whether Asn promotes bone and lung metastases by acting as a chemoattractant during metastatic dissemination. To this end, we performed Boyden chamber in the presence or absence of an Asn gradient (0.1 mM in the lower chamber, approximating physiological tissue concentrations^11,15^. Asn-containing medium did not enhance cell migration (Suppl. Fig. 1H, I), indicating that Asn does not function as a chemotactic cue guiding metastatic dissemination in PC.

We therefore hypothesized that elevated Asn availability facilitates metastasis colonization once DTCs reach and begin to colonize the secondary site. To test this hypothesis, we employed three-dimensional spheroid cultures (3D-C), an experimental model widely used to mimic the metabolic adaptations acquired by DTCs along metastatic dissemination^9,16–18^. Depletion of low-molecular-weight components from FBS (dFBS, dialysis cutoff ≤3.5 kDa) markedly impaired 3D-C formation (evaluated as the average area of 3D-C clusters) (Suppl. Fig. 1J), indicating that small metabolites, including amino acids, contribute substantially to this process. Notably, supplementation of Asn (0.1 mM) in the dFBS-containing medium was sufficient to restore and enhance the area of 3D-C (in two different metastatic cell lines PC3 and DU-145) (Fig. 1H, I), while no effect on 2D cell expansion was observed (Suppl. Fig. 1 K, L). In keeping, depletion of extracellular Asn using L-asparaginase (ASNase, Suppl. Fig. 1M) abolished the Asn-dependent increase in 3D-C area (Fig. 1J). In addition to its asparaginase activity, ASNase also exhibits glutaminase activity. Consistent with this, ASNase treatment in our model reduced extracellular glutamine (Gln) levels even in the presence of Asn (Suppl. Fig. 1N). To exclude the possibility that the observed effects were attributable to glutamine depletion rather than Asn degradation, we cultured 3D-C under Gln-deprived conditions. Under Gln restriction 3D-C area was not affected, and the advantage conferred by Asn supplementation was preserved (Suppl. Fig. 1O), indicating that the effects of ASNase are independent on its glutaminase activity. These findings support the idea that elevated Asn availability represents an advantage for DTCs during niche colonization. To generate a tissue-specific *ex-vivo* model of early metastatic PC cells, we injected mice with GFP⁺ PC3 cells either i.c. or *via* the caudal artery (c.a.)^19^. At 9 weeks post-injection (a time point consistent with early stages of metastatic formation in *in vivo* PC models), tibial bones, lungs, and liver were harvested, mechanically dissociated, and cancer cells were purified by FACS based on GFP fluorescence (Fig. 1K, Suppl. Fig. 2A). Notably, Asn supplementation enhanced the size of bone- (B-M-1, B-M-2) (Fig. 1L, M) and lung-derived (L-M-1, L-M-2) (Fig. 1N-O) metastatic cells 3D-C spheroids, while liver-derived (Liv-M) (Fig. 1P) ones were not affected.

Together, these findings reveal that bone- and lung-specific enrichment of environmental Asn supports incoming PC DTCs in metastatic colonization.

### 2. Exogenous Asn increase 3D-C spheroids formation by sustaining enhanced translational program

Asn is a non-essential amino acid that can be taken up from the extracellular environment or synthesized *de novo* by asparagine synthetase (ASNS), which catalyzes the conversion of Asp to Asn using Gln as a nitrogen donor. Loss of cell adhesion has previously been associated with reduced expression of activating transcription factor 4 (ATF4) and its downstream targets, including ASNS^18^. Consistently, ATF4 and ASNS expression was reduced in 3D-C cells compared with 2D conditions (Fig. 2A-B) suggesting that tissue-incoming DTCs mostly depend on extracellular Asn rather than activating the endogenous biosynthetic pathway. A similar downregulation was detected in lung- and bone-derived metastatic cells compared with parental counterparts (wt GFP⁺ PC3, prior to mice injection) (Suppl. Fig. 3A). In keeping, supplementation with physiological levels of Asp (0.15 mM) failed to enhance 3D-C area (Fig. 2C), indicating that Asn provides an advantage independently of Asp-to-Asn conversion.

**Fig. 2.**
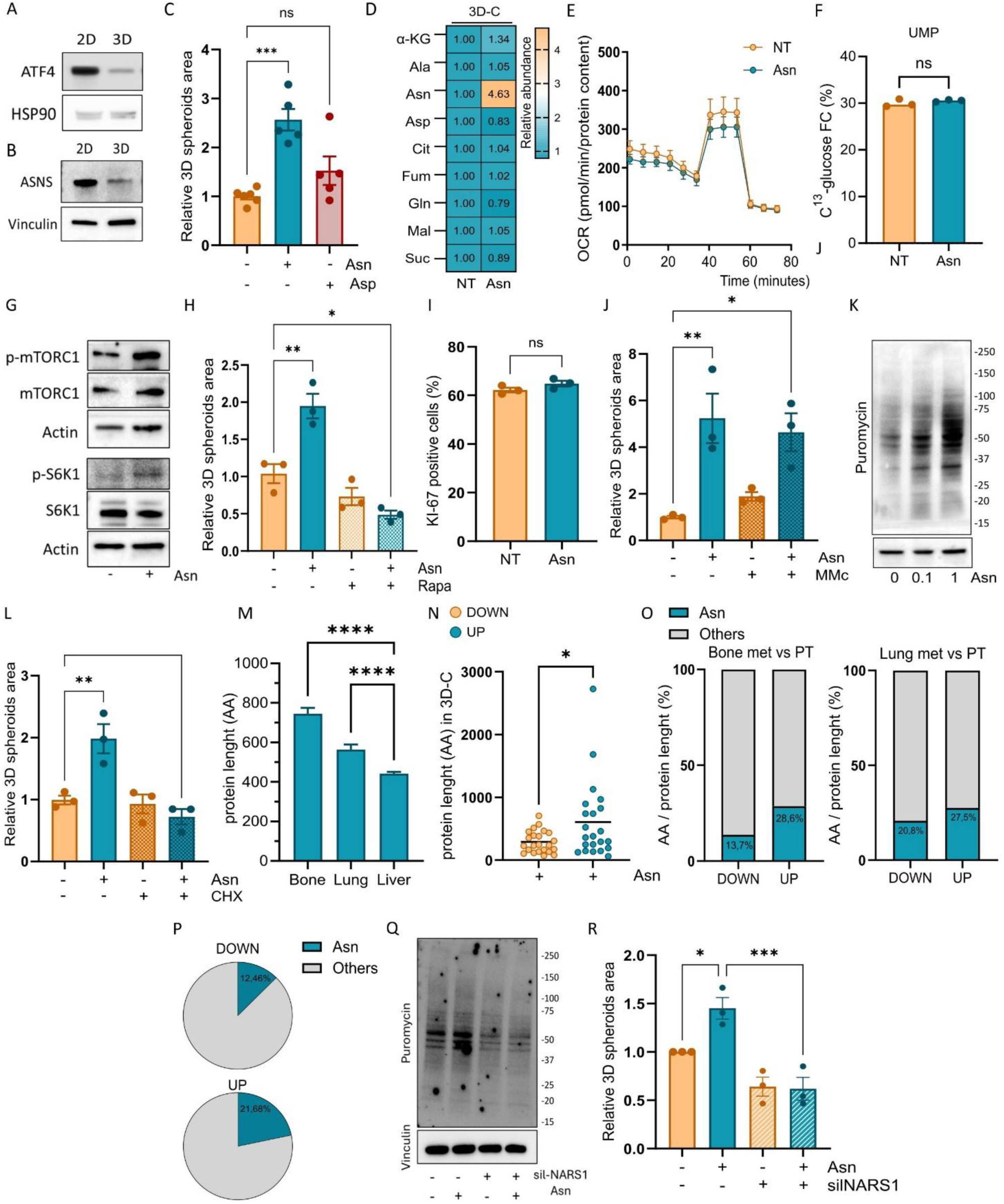
PC cells cultured in 3D-C exhibit reduced ASNS expression and depend on exogenous Asn to activate mTORC1 signaling and increase protein synthesis. **(A, B)** Western blot analysis of ATF4 (A) and ASNS (B) in PC3 cells cultured in standard two-dimensional (2D) conditions or in 3D-C. HSP90 (A) and vinculin (B) were used as loading controls. Images are representative of three independent experiments. **(C)** Total spheroid area of PC3 3D-C grown with or without Asn (0.1 mM) in the presence or absence of aspartate (Asp, 0.15 mM). One-way ANOVA with Sidak’s correction. **(D)** Relative abundance of Asn-related metabolites in PC3 3D-C cultured with or without Asn (0.1 mM) for 5 days, quantified by GC-MS. Valued are reported as relative to non-treated control (NT). No major changes were observed except for a selective increase in intracellular Asn. Reported values are average of three independent experiments. **(E)** Oxygen consumption rate (OCR) evaluated by Seahorse analysis in PC3 3D-C treated or not with Asn (0.1 mM) for 5 days. Values are reported as relative to non-treated (NT) control. **(F)** Fractional enrichment of ^13^C-UMP after 24 h labeling with U-^13^C-glucose in PC3 3D-C treated or not with Asn (0.1 mM) for 5 days. Welch’s t-test. **(G)** Phosphorylation of mTORC1 and its downstream target p70S6K in PC3 3D-C cultured with or without Asn (0.1 mM) for 5 days. Data are normalized to total mTORC1 or total p70S6K and actin. Images are representative of three independent experiments. **(H)** Total spheroid area of PC3 3D-C grown with or without Asn (0.1 mM) in the presence or absence of rapamycin (Rapa, 0.01 mM) for 5 days. One-way ANOVA with Dunnett’s correction. **(I)** Quantification of Ki-67⁺ proliferating cells in PC3 3D-C cultured with or without Asn (0.1 mM), expressed as percentage of total cells and measured by flow cytometry. Unpaired Student’s t-test. **(J)** Total spheroid area of PC3 3D-C grown with or without Asn (0.1 mM) in the presence or absence of mitomycin C (MMc, 1 mM). One-way ANOVA with Dunnett’s correction. **(K)** Total protein synthesis in PC3 3D-C cultured with or without Asn (0.1 or 1 mM) for 5 days, assessed by puromycin incorporation as in panel A. Actin was used as a loading control. The image is representative of three independent experiments. **(L)** Total spheroid area of PC3 3D-C grown with or without Asn (0.1 mM) in the presence or absence of cycloheximide (CHX, 0.1 mM). One-way ANOVA with Dunnett’s correction. ns, not significant; **(M)** Average protein length (number of amino acids) encoded by genes upregulated in bone-, lung-, and liver-derived metastatic cells isolated from NOD-SCID mice injected intracardially (i.c.) with GFP⁺ PC3 cells. Metastatic lesions were collected 9–12 weeks post-injection, dissociated, and GFP⁺ cancer cells were isolated by FACS for RNA sequencing. One-way ANOVA with Sidak’s correction. **(N)** Average length of proteins (number of amino acids) significantly upregulated or downregulated in PC3 3D-C cultured in presence of Asn (0.1 mM) for 5 days. Proteins specific to 3D-C were obtained by removing proteins also expressed in 2D conditions. Welch’s t-test. **(O)** Asn residue content in proteins encoded by genes significantly up- or downregulated in bone- and lung-derived metastatic cells relative to the primary tumor (PT). RNA-seq analysis was conducted as described in (M). Values are expressed as percentage of total amino acid residues. **(P)** Relative Asn residue content in proteins significantly upregulated or downregulated in PC3 3D-C treated with Asn (0.1 mM) compared to untreated 3D-C. Values are expressed as percentage of total amino acid residues. **(Q)** Total protein synthesis in PC3 cells silenced for NARS1 for 48 h prior to 3D-C culture with or without Asn (0.1 mM), assessed by puromycin incorporation as in panel A. Vinculin was used as a loading control. The image is representative of three independent experiments. **(R)** Total spheroid area of NARS1-silenced PC3 cells grown in 3D-C with or without Asn (0.1 mM). One-way ANOVA with Dunnett’s correction. *P < 0.05; **P < 0.01; ***P < 0.001. Data represent mean ± s.e.m. from at least three independent experiments.

We next investigated whether alternative metabolic functions of Asn support 3D-C metabolism. Although Asn has been implicated in maintaining amino acid homeostasis under nutrient deprivation, particularly Gln^20^, in regulating mitochondrial respiration^21^, and in sustaining nucleotide biosynthesis^22^, its supplementation in 3D-C did not alter the steady-state levels of metabolites associated to Asn and Asp metabolism assessed by GC-MS analysis (Fig. 2D). Similarly, Seahorse analysis revealed no significant changes in oxygen consumption rate in 3D-C upon Asn supplementation (Fig. 2E). In addition, ^13^C-glucose tracing did not detect increased pyrimidine nucleotide biosynthesis in Asn-treated 3D-C, as indicated by unchanged fractional incorporation of glucose-derived carbons into uridine monophosphate (UMP) (Fig. 2F). Together, these data indicate that Asn does not affect mitochondrial respiration or pyrimidine nucleotide synthesis in this context.

Elevated extracellular Asn has been also reported to activate mTORC1 pathway^23^ leading to phosphorylation of its downstream effector ribosomal protein S6 kinase beta-1 (S6K1)^24^, a key regulator of protein synthesis^25^. Consistent with this, Asn supplementation in 3D-C triggered mTORC1 activation and increased S6K1 phosphorylation (Fig. 2G). In keeping with a functional role of Asn-mediated mTORC1 activation, the treatment with the mTORC1 inhibitor Rapamycin abolished the Asn-mediated increase in 3D-C area (Fig. 2H). Activated mTORC1 pathway is known to sustain cell proliferation and protein synthesis^26^. First, we excluded the contribution of extracellular Asn to cell proliferation in 3D-C, as Ki-67 staining was unchanged (Fig. 2I), and pharmacological inhibition of proliferation with mitomycin C failed to abrogate the Asn-induced increase in 3D-C area (Fig. 2J). On the other hand, we observed a dose-dependent Asn-driven increase in intracellular protein synthesis in 3D-C, as demonstrated by the SUnSET analysis (Fig. 2K). Consistently, pharmacological inhibition of protein synthesis with Ciclohexamide abrogated the increase in 3D-C area conferred by Asn (Fig. 2L). Together, these data demonstrate that Asn sustains DTCs translational program.

### 3. Bone and lung metastatic cells rewire translation toward Asn-rich proteins during early metastatic colonization

Metastatic colonization requires extensive remodeling of the cellular proteome^27,28^. We therefore hypothesized that elevated Asn availability does not globally enhance protein synthesis, but rather selectively favors the production of specific subsets of proteins required for the colonization of novel ECM environments (i.e. bone and lung). These include membrane-associated and extracellular proteins^29^, which are typically characterized by long amino acid sequences and extensive post-translational modifications. We started by analyzing transcriptomic profiles of RNA freshly extracted from FACS-sorted PC cells isolated from primary tumors and cancer cells collected from bone, lung, and liver of mice previously injected with GFP⁺ PC3 cells. RNA-seq analysis revealed that bone- and lung-derived metastatic cells cluster separately from primary tumors and liver metastases (Suppl. Fig. 2B). Specifically, genes upregulated in bone- and lung-derived metastatic cells encoded proteins with greater amino acid length compared with those enriched in liver-derived metastases, suggesting an increased demand for the synthesis of longer proteins in cells colonizing bone and lung (Fig. 2M). Notably, this feature was recapitulated *in vitro* by mimicking the elevated Asn availability encountered by DTCs upon arrival at bone and lung niches. Indeed, proteomic profiling of 3D-C exposed or not to Asn showed that upregulated proteins upon Asn supplementation were, on average, significantly longer than downregulated ones (Fig. 2N, Suppl. Tab1-2).

Because the availability of specific amino acids can influence tRNA charging and modulate translation rates of specific subsets of proteins^30^, we next asked whether proteins upregulated upon Asn exposure were specifically enriched in Asn residues. Notably, proteins encoded by genes upregulated in bone- and lung-derived metastatic cells were characterized by a higher proportion of Asn residues compared with those that were downregulated (Fig. 2O). Consistently, proteome analysis revealed that also proteins upregulated upon Asn supplementation in 3D-C were characterized by a higher Asn content than downregulated ones (Fig. 2P). To directly assess the functional requirement for Asn incorporation into nascent polypeptides, we silenced asparaginyl-tRNA synthetase 1 (NARS1), the enzyme responsible for charging tRNA with Asn (Suppl. Fig.3B-C). NARS1 silencing markedly reduced global protein synthesis in 3D-C (Fig. 2Q) and abrogated the Asn-mediated increase in 3D-C spheroid formation (Fig. 2R) while having minimal effect on cells in 2D culture (Suppl. Fig. 3D). Collectively, these results indicate that elevated availability of Asn in bone and lung supports the synthesis of Asn-rich proteins in PC metastasizing cells.

**Fig. 3.**
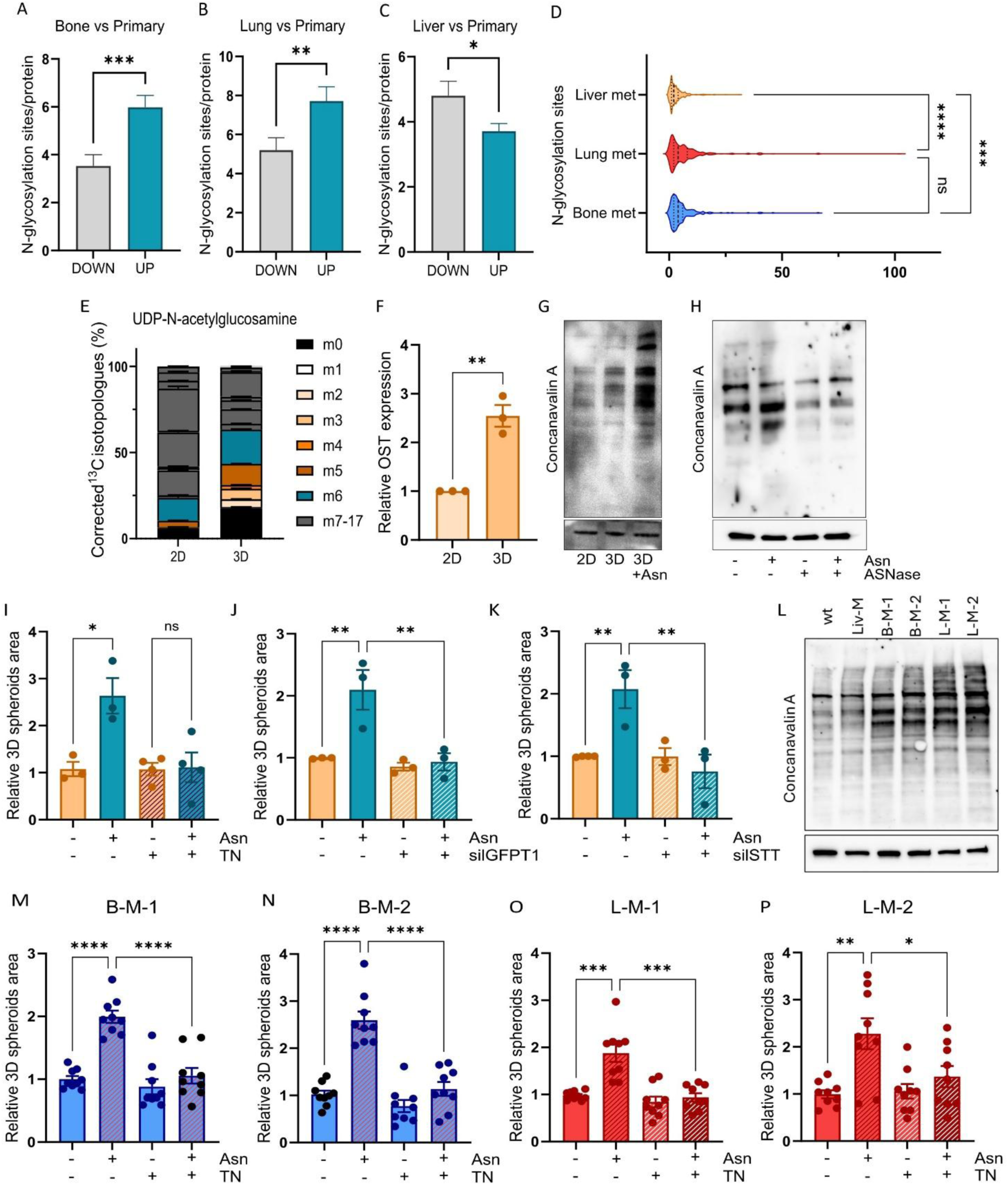
Enhanced protein N-glycosylation underlies Asn-driven increase in 3D-C growth. **(A-C)** Number of putative N-glycosylation sites in proteins encoded by genes upregulated in bone-(A), lung-(B), and liver-(C) derived metastatic cells relative to the primary tumor (PT). RNA-seq analysis was conducted as described in (Fig. 2N). Values are expressed as relative to proteins number. Welch’s t-test. **(D)** Number of putative N-glycosylation sites in proteins encoded by genes upregulated in bone-, lung-, and liver-derived metastatic cells. One-way ANOVA with Tukey’s correction. **(E)** Relative incorporation of U-^13^C-glucose carbons in UDP-N-acetyl-glucosamine. PC3 cells were grown in 3D-C for 5 days and subsequently incubated in a medium containing U-^13^C-glucose for 24h. Metabolite abundance and labeling enrichment were evaluated by LC-MS analysis. **(F)** mRNA levels of oligosaccharyltransferase (OST) in 2D coltures and 3D-C. mRNA expression levels analyzed by quantitative RT-PCR using 2D condition as comparator. Student’s t test. **(G)** Concanavalin A lectin binding assay performed on lysates from PC3 cells cultured in 2D conditions or 3D-C, with or without Asn supplementation (0.1 mM). Immunoblot for actin was used to confirm equal protein loading across samples. The image is representative of three independent experiments. **(H)** Concanavalin A lectin binding assay performed on lysates from 3D-C cultured with or without Asn (0.1 mM) in the presence or absence of L-asparaginase (ASNase, 0.25 U/ml). Immunoblot for vinculin was used to confirm equal protein loading across samples. The image is representative of three independent experiments. **(I)** Total spheroid area of PC3 3D-C grown with or without Asn (0.1 mM) in the presence or absence of Tunicamycin (5ng/µl). One-way ANOVA with Dunnett’s correction. **(J)** Total spheroid area of Glutamine-fructose-6-phosphate transaminase 1 (GFPT1)-silenced PC3 cells grown in 3D-C with or without Asn (0.1 mM). One-way ANOVA with Dunnett’s correction. **(K)** Total spheroid area of 3D-C PC3 cells silenced for the OST catalytic subunits STT3A and STT3B with or without Asn (0.1 mM). One-way ANOVA with Dunnett’s correction. **(L)** Concanavalin A lectin binding assay performed on lysates from liver-derived (Liv-M), bone-derived (B-M-1, B-M-2) and lung-derived (L-M-1, L-M-2) metastatic cells (obtained after intracardiac or caudal artery injection, respectively) grown in 3D-C cultures. Immunoblot for vinculin was used to confirm equal protein loading across samples. The image is representative of three independent experiments. **(M-P)** Total spheroid area of bone-derived (M, N), and lung-derived (O, P) metastatic cells grown in 3D-C cultures with or without Asn (0.1 mM) in the presence or absence of Tunicamycin (5ng/µl). Two-way ANOVA with Tukey’s correction. ns, not significant; *P < 0.05; **P < 0.01; ***P < 0.001; ****P < 0.0001. Data represent mean ± s.e.m. from at least three independent experiments.

### 4. Exogenous Asn is exploited to sustain N-glycosylated protein synthesis in bone- and lung-colonizing PC cells

Asn accounts for approximately 4-5% of amino acid residues within human proteins, placing it not among the most represented amino acids^31^. Nonetheless, Asn is unique as it is the sole amino acid residue within the consensus motif Asn-X-Ser/Thr that undergoes N-linked glycosylation, a co-translational modification that occurs in the endoplasmic reticulum (ER). N-glycosylation is essential for correct protein folding and stability, and critically regulates protein trafficking, secretion, and functional activity, particularly for membrane-bound and extracellular proteins^32,33^. Based on these considerations, we hypothesized that Asn availability may preferentially support the synthesis of proteins destined to N-glycosylation process. Interestingly, transcriptomic analysis of bone-, lung-, and liver-derived metastatic cells revealed that proteins encoded by genes upregulated in bone- and lung-derived metastases (but not in liver-derived ones) were significantly enriched in predicted N-glycosylation sites compared to proteins upregulated in primary tumors (Fig. 3A-C; Suppl. Fig. 3E-G). Moreover, the number of predicted N-glycosylation sites per proteins was significantly higher in bone- and lung-derived metastases than in liver-derived ones (Fig. 3D).

The hexosamine biosynthetic pathway (HBP) is a metabolic pathway that consumes approximately 2–5% of cellular glucose to generate UDP-N-acetylglucosamine (UDP-GlcNAc), which is the principal substrate used for N-linked glycosylation of proteins^34^. We therefore investigated differential HBP activity using ^13^C-glucose tracing in 2D cultures and 3D-C. The expected mass shift of UDP-GlcNAc is M+17, comprising M+6 from the glucose moiety incorporated and converted in fructose 6-phosphate (the HBP precursor), M+2 from the acetyl moiety derived from acetyl-CoA via glycolysis followed by PDH complex activity, M+5 from the ribose moiety via the pentose phosphate pathway (PPP), and for long-term incubations (i.e. 24h), M+4 from uracil derived from aspartate via TCA cycle^35,36^. Interestingly, we observed a shift in fractional labeling in the 3D-C (Fig. 3E) consistent with increased glucose routing through HBP (increased M+6, Suppl. Fig.3H) and contribution of ribose from PPP (increased M+5, Suppl. Fig.3I) compared to 2D cultures. Conversely, quantification of pentose isotopologues revealed a reduced fraction of fully labeled (M+5) species in 3D-C compared with 2D conditions, indicating a decreased flux of glucose to pentose phosphate pathway and consequently nucleotide biosynthesis (Suppl. Fig. 3J). N-linked glycosylation is catalyzed in the ER by the oligosaccharyltransferase (OST) complex, which transfers a preassembled oligosaccharide from the lipid carrier dolichol to specific Asn residues within the consensus sequence of nascent polypeptides. Consistent with previous data, OST expression was upregulated in 3D-C compared with 2D cultures (Fig. 3F). Together, these data indicate that cells in 3D-C are metabolically predisposed to increase protein N-glycosylation. We therefore hypothesize that, when Asn availability is sufficient, this may result in enhanced synthesis of N-glycoproteins. Consistently, Concanavalin A blot analysis, which detects N-glycosylated proteins through specific binding to mannose- and glucose-containing glycans, revealed an increase in global protein N-glycosylation in 3D-C compared to 2D cultures, with a markedly stronger enhancement in 3D-C upon Asn supplementation (Fig. 3G). Conversely, depletion of extracellular Asn using ASNase abolished the Asn-induced increase in protein N-glycosylation in 3D-C (Fig. 3H). Accordingly, pharmacological inhibition of N-linked glycosylation with tunicamycin, which specifically targets UDP-N-acetylglucosamine: dolichol phosphate N-acetylglucosamine-1-phosphate transferase (GPT), the enzyme catalyzing the first committed step of N-linked glycosylation, abolished the Asn-driven advantage in 3D-C (Fig. 3I). Similarly, silencing the HBP rate-limiting enzyme glutamine-fructose-6-phosphate transaminase 1 (GFPT1) (Suppl. Fig. 3K) or the OST complex *via* co-silencing of STT3A and STT3B (Suppl. Fig. 3L) decreased total protein N-glycosylation (Suppl. Fig. 3M) and abolished the Asn-mediated increase in 3D-C area (Fig. 3J-K). Notably, we observed a marked increase in protein N-glycosylation in B-M-1/B-M-2 and L-M-1/L-M-2 cells compared with the parental counterparts, whereas no increase was detected in Liv-M cells (Fig. 3L). Consistently, Tunicamycin treatment impaired the Asn-mediated increase in 3D-C area in bone- and lung-derived metastatic lines (Fig. 3M-P). Together, these findings indicate that Asn preferentially supports the synthesis of proteins destined for N-glycosylation in bone- and lung-colonizing DTCs.

### 5. Asn promotes N-glycosylation-driven cell-cell and cell-matrix adhesion in PC metastatic cells

Pathway enrichment analysis of proteomes of 3D-C exposed or not to Asn revealed increased expression of proteins involved in translation and enhanced trafficking toward the endoplasmic reticulum (ER), consistent with an elevated demand for secretory and membrane proteins (Fig. 4A). Given the central role of secretory and surface proteins in mediating intercellular interactions, we hypothesized that the enhanced area of 3D-C spheroids upon Asn supplementation resulted from increased cell-cell adhesion. Supporting this hypothesis, EDTA treatment (used to disrupt cadherin-mediated adhesion) completely abolished the advantage conferred by Asn (Fig. 4B). Consistently, in a short-term aggregation assay (6 h incubation on non-adherent plates) (Fig. 4C), cells dissociated from already established 3D-C (after 5 days of suspension culture) exhibited a markedly greater capacity to re-aggregate than cells exposed to non-adherent condition for the first time (i.e., directly transferred from 2D cultures) (Fig. 4D). Culturing 3D-C for 5 days in the presence of Asn further enhanced their re-aggregation capacity, whereas Asn supplementation restricted to the short-term aggregation assay (6h), without prior administration during 3D-C growth, failed to promote cell clustering (Fig. 4E). In line with these findings, depletion of extracellular Asn using ASNase during the 5 days culture period abolished the Asn-dependent aggregation advantage (Fig. 4F). Importantly, impairing Asn incorporation into nascent proteins by silencing NARS1 eliminated the re-aggregation ability of 3D-C-derived cells, independently of Asn availability (Fig. 4G). Collectively, these findings indicate that high Asn availability sustains the synthesis of a subset of Asn-rich proteins, likely involved in cell-cell adhesion.

**Fig. 4.**
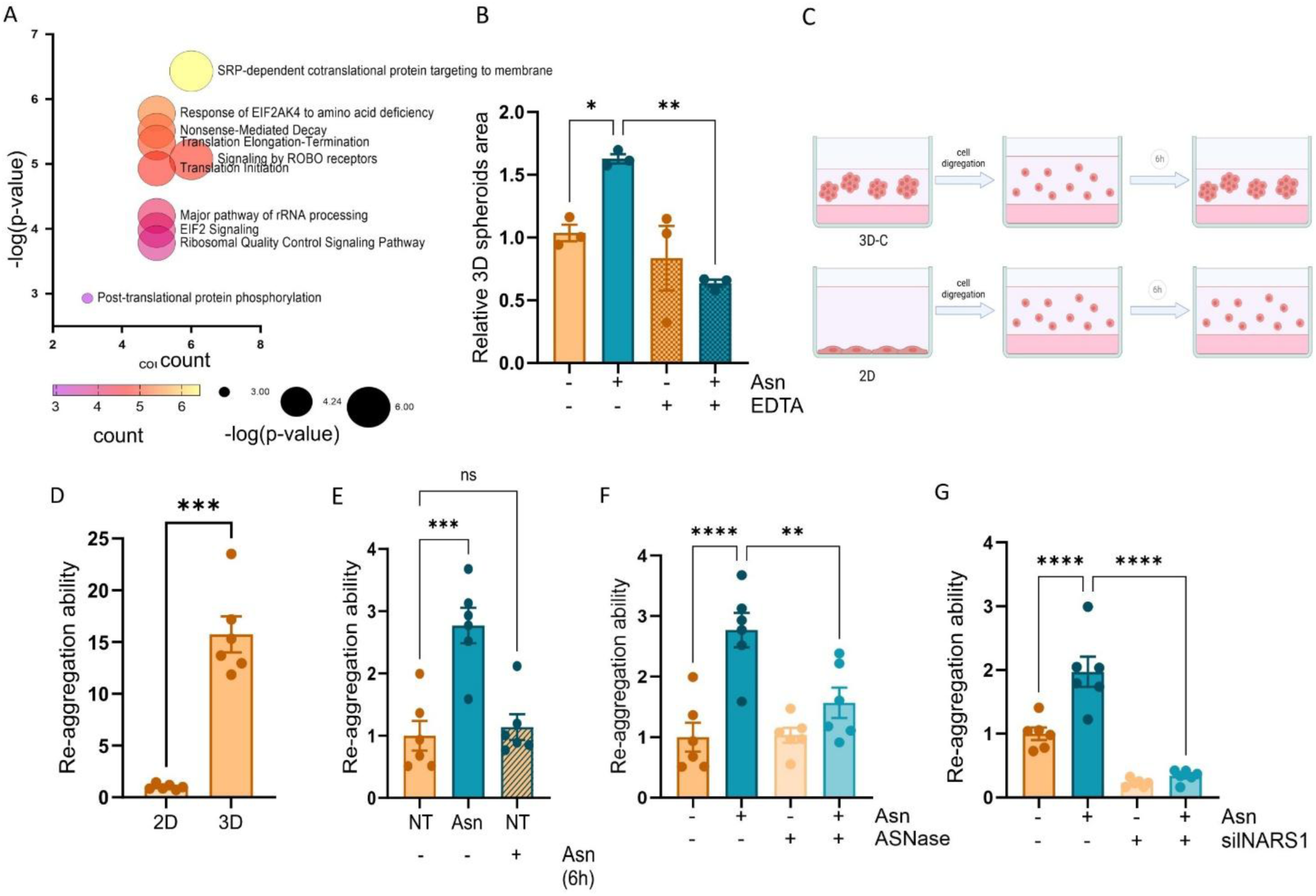
Exogenous Asn sustains metastatic colonization supporting the synthesis of proteins involved in cell-cell adhesion. **(A)** Pathway enrichment analysis of proteomic data comparing PC3 3D-C cultured with Asn (0.1 mM) versus untreated 3D-C. Proteins specific to 3D-C were obtained by excluding those also expressed in 2D conditions. The color scale represents −log10 adjusted P value (Y-axis). Circle size (X-axis) indicates the number of significant proteins associated with each biological process. **(B)** Total spheroid area of PC3 3D-C grown with or without Asn (0.1 mM) in the presence or absence of EDTA (0.1 mM). One-way ANOVA with Dunnett’s correction. **(C)** Schematic representation of short-term aggregation assays (D-G). Cells from 5 days of incubation in either 3D-C (with or without Asn 0,1 mM) or normal 2D cultures were dissociated and replated onto non-adherent plates for 6h. For the experiment shown in Fig. 4E, acute Asn (0,1 mM) supplementation was applied during the aggregation assay (6h). **(D-G)** Total area of PC3 cell clusters aggregated during 6 h of incubation onto agar-coated non-adherent plates. (D) Comparison between cells derived from conventional 2D cultures and established 3D-C; **(E)** Effect of Asn supplementation during 3D-C growth (5-days of incubation) versus short-time Asn incubation (6h, limited to the aggregation assay); **(F)** Effect of extracellular Asn depletion by L-asparaginase (ASNase, 0.25 U/ml) during the 5-day 3D-C culture period; **(G)** Effect of NARS1 silencing in PC3 cells during the 5-day 3D-C culture period in the presence or absence of Asn. One-way ANOVA with Sidak’s correction. ns, not significant; *P < 0.05; **P < 0.01; ***P < 0.001; ****P < 0.0001. Data represent mean ± s.e.m. from at least three independent experiments.

Together with cell-cell adhesion, cell-ECM interaction is also essential for the successful metastatic seeding and colonization. Interestingly, REACTOME pathway analysis of transcriptomic profiles of parental cells (wt GFP⁺ PC3) and bone-/lung-associated metastatic lesions revealed that bone- and lung-derived metastasis upregulate programs associated with ECM organization and interaction as well as altered protein glycosylation, suggesting that PC metastasizing cells enhance connections with the bone and lung microenvironment (Fig. 5A-B). To assess the contribution of Asn to cell-ECM interactions, we performed matrix-adhesion assays: 3D-C were allowed to adhere to native decellularized extracellular matrices derived from non-diseased porcine bone, lung, or liver (Fig. 5C). Asn supplementation significantly increased the adhesion to bone- and lung-derived matrices (Fig. 5D-E), but not to the liver-derived one (Fig. 5F). Notably, bone and lung ECMs share a high fibrillar collagen I and hyaluronic acid content^37–40^, whereas the liver ECM is predominantly composed of a reticular/basement membrane-type matrix enriched in collagen IV and laminins^41,42^. These structural differences may underlie the selective advantage conferred by Asn in bone and lung metastases. In keeping, similar results were obtained using individual ECM components abundant in bone and lung, such as collagen type I (Suppl. Fig. 3N) and hyaluronic acid (Suppl. Fig. 3O). Depletion of extracellular Asn with ASNase abolished the Asn-dependent increase in the adhesion to collagen I (Fig. 5G). Notably, providing Asn exclusively during the adhesion assay (without prior exposure during 3D-C growth) failed to enhance the adhesion to collagen type I (Fig. 5H) compared to the Asn-experienced 3D-C, suggesting that Asn availability is required to induce long-term adaptations such as protein synthesis and modifications (i.e N-glycosylation), rather than acute signaling effects. In line with this, preventing Asn incorporation into nascent proteins through NARS1 silencing (Fig. 5I) as well as the pharmacological (Fig. 5J)/genetic (Fig. 5K, L) inhibition of N-linked glycosylation abolished the Asn-mediated increase in the adhesion to collagen I. Together, these results indicate that Asn promotes the adhesion to bone/lung ECM by sustaining the synthesis of N-linked glycoproteins.

**Fig. 5.**
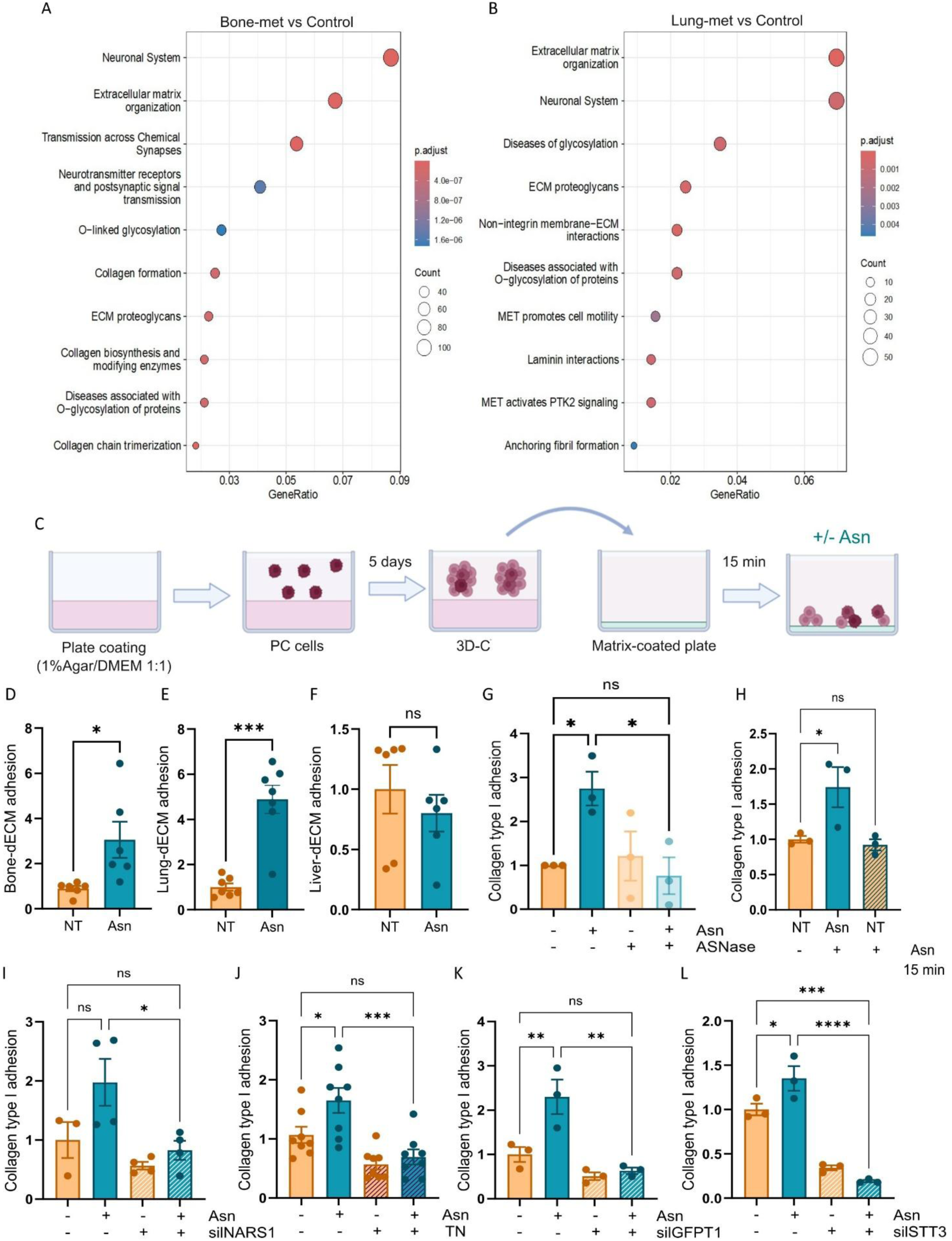
Asn-dependent N-glycoprotein synthesis is required for bone/lung niche colonization by enhancing cell-matrix adhesion. **(A-B)** Pathway enrichment analysis of RNA-seq data. Dot plots show Reactome pathways enriched among upregulated genes in metastatic cells isolated from bone (A) and lung (B) tissues 12 weeks after intracardiac (i.c.) injection, compared with control PC3 cells (top 10 upregulated Reactome gene sets). The x-axis indicates the gene ratio for each pathway, dot size reflects the number of genes annotated to each pathway, and dot color represents the adjusted p value (red denotes higher significance). **(C)** Schematic representation of the adhesion assay used to evaluate the ability of cells cultured for 5 days in 3D-C to adhere to matrix-coated plates. For the experiment shown in Fig. 4H, acute Asn (0.1 mM) supplementation was applied during the adhesion assay. **(D-F)** Adhesion to decellularized extracellular matrices. PC3 3D-C were cultured with or without Asn (0,1 mM) for 5 days and allowed to adhere for 15 min to plates coated with decellularized matrices derived from non-diseased porcine bone (D), lung (E), or liver (F), followed by quantification of adherent cells. Data are shown relative to untreated cells. Welch’s t test. **(G-L)** Collagen type I adhesion assays. (G) Adhesion assay on 3D-C cultured with or without Asn (0.1 mM) in the presence or absence of L-asparaginase (ASNase, 0.25 U/ml) for 5 days before adhesion assay. One-way ANOVA with Sidak’s correction. (H) Effect of Asn supplementation during 3D-C growth (5-days of incubation) versus short-time Asn incubation (15 min, limited to the adhesion assay). One-way ANOVA with Dunnett’s correction. (I) Adhesion assay on PC3 cells silenced for asparaginyl-tRNA synthetase 1 (NARS1) prior to establishing 3D-C with or without Asn (0.1 mM). One-way ANOVA with Sidak’s correction. (J) Adhesion assay on PC3 3D-C grown with or without Asn (0.1 mM) in the presence or absence of Tunicamycin (5ng/µl). One-way ANOVA with Sidak’s correction. (K) Adhesion assay on Glutamine-fructose-6-phosphate transaminase 1 (GFPT1)-silenced PC3 cells grown in 3D-C with or without Asn (0.1 mM). One-way ANOVA with Dunnett’s correction. (L) Adhesion assay on 3D-C PC3 cells silenced for the OST catalytic subunits STT3Aand STT3B with or without Asn (0.1 mM). One-way ANOVA with Dunnett’s correction. ns, not significant; *P < 0.05; **P < 0.01; ***P < 0.001; ****P < 0.0001. Data represent mean ± s.e.m. from at least three independent experiments.

### 6. CD44 upregulation drives Asn-mediated cell–cell and cell–matrix interactions

To identify specific effectors of Asn-mediated metastatic adaptation, we prioritized membrane-associated proteins among the top upregulated candidates identified by proteomic analysis (≥1.5-fold, Asn 3D-C vs NT-3D-C, 2D-filtered) (Fig. 6A, Suppl. Tab.3). This approach allowed us to identify CD44 as a valuable target for further investigations. CD44 is a heavily N-glycosylated adhesion receptor^43^ that mediates binding to hyaluronic acid. Consistently, B-M-1/B-M-2 and L-M-1/L-M-2 cells exhibited higher CD44 protein levels compared to parental counterparts (Fig. 6B). Western blot analysis further showed that CD44 expression was higher in 3D-C than in 2D cultures (Fig. 6C), and progressively increased during 3D-C formation (Fig. 6D). Supporting the clinical relevance of these findings, IHC analysis of PC patient samples revealed higher CD44 expression in bone metastases compared with primary tumors (Fig. 6E, F). The functional relevance of CD44 was confirmed by gene silencing (Suppl. Fig. 3P), which impaired cell-cell adhesion (Fig. 6G) and hyaluronic acid adhesion (Fig. 6H) in 3D-C.

**Fig. 6.**
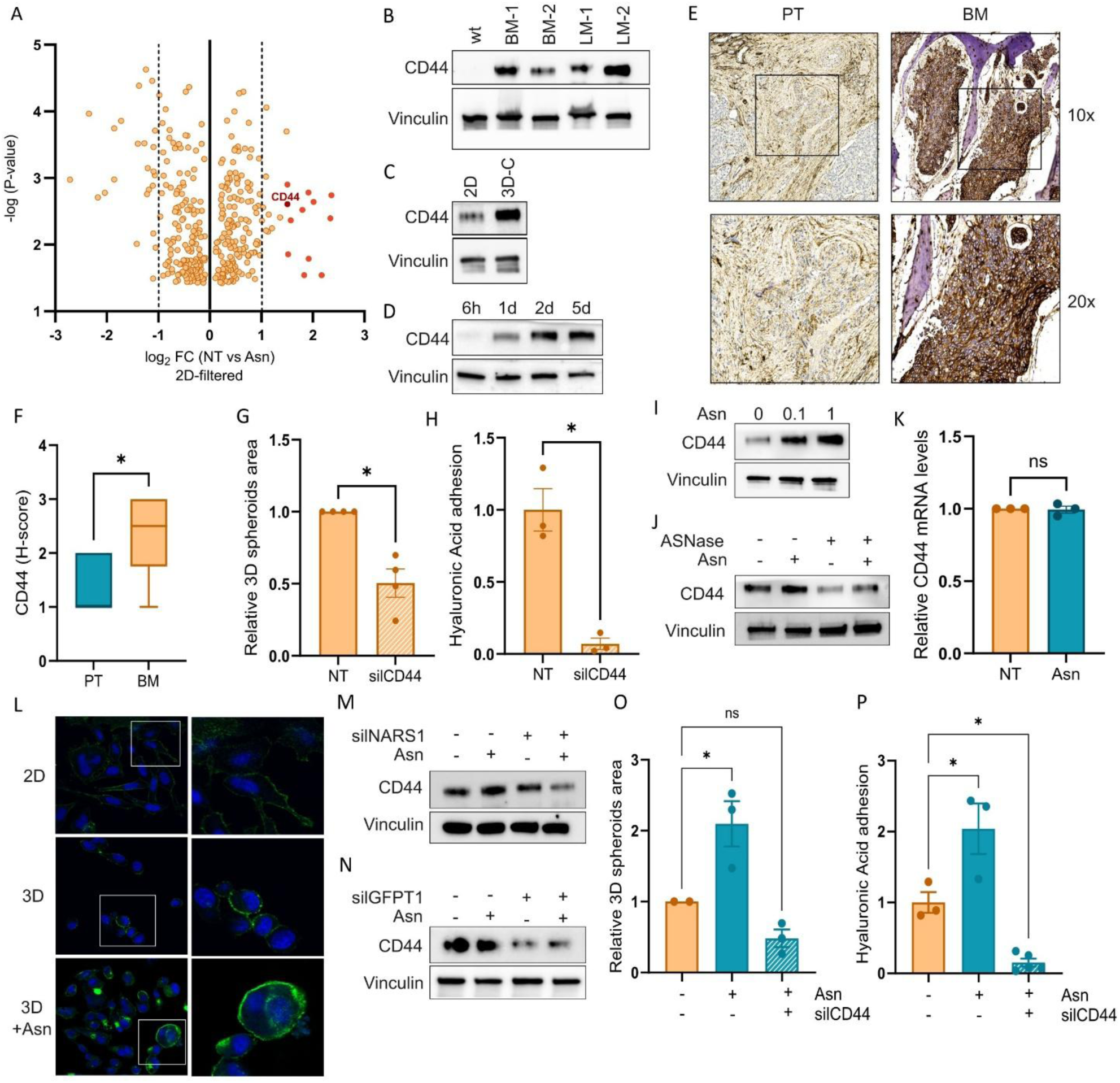
Asn-driven CD44 upregulation promotes cell-cell and cell-matrix interactions. **(A)** Volcano plot showing differential protein abundance in PC3 cells grown in 3D-C in the absence or presence of Asn (0.1 mM). The x-axis reports the difference (log_2_ fold change) fold change between 3D-C and 3D-C+Asn; the y-axis shows –log_10_(p value). Proteins specific to 3D-C were identified by excluding proteins also expressed in 2D conditions. Each dot represents a quantified protein; red dots denote proteins exceeding the ≥2-fold change threshold. **(B)** Western blot analysis of CD44 expression in metastatic cells isolated from bone (B-M-1, B-M-2) and lung (L-M-1, L-M-2) lesions isolated as described in Fig. 1K. Vinculin was used as a loading control. Representative of three independent experiments. **(C)** Western blot analysis of CD44 in PC3 cells cultured under standard 2D conditions or in 3D-C. Vinculin was used as a loading control. Representative of three independent experiments. **(D)** Time-course Western blot analysis of CD44 expression in PC3 cells cultured in 3D-C during spheroid formation (5 h, 1 day, 2 days, and 5 days after plating). Vinculin was used as a loading control. The image is representative of three independent experiments. **(E)** Representative immunohistochemical staining for CD44 in primary prostate carcinoma tissues (PT) and bone metastatic (BM) lesions from patients with PC. Boxed areas indicate regions shown at higher magnification. Brown staining denotes CD44-positive cells; nuclei are stained with hematoxylin. **(F)** H-score quantification of CD44 immunohistochemical staining shown in (E). H-scores were calculated by integrating staining intensity and the percentage of positive cells. Welch’s *t* test. **(G)** Total spheroid area of CD44-silenced PC3 cells grown in 3D-C compared to control silencing condition. Welch’s t-test. **(H)** Hyaluronic acid adhesion assay on CD44-silenced PC3 cells grown in 3D-C for 5days before plating on hyaluronic acid-coated plate. Data are shown relative to untreated cells. Welch’s t test. **(I)** Western blot analysis of CD44 expression in PC3 cells cultured in 3D-C in the presence or absence of Asn (0.1 mM and 1 mM). Vinculin was used as a loading control. The image is representative of three independent experiments. **(J)** Western blot analysis of CD44 in PC3 cells in 3D-C with or without Asn (0.1 mM) in the presence or absence of L-asparaginase (ASNase, 0.25 U/ml). Vinculin was used as loading control. The image is representative of three independent experiments. **(K)** CD44 mRNA expression levels in PC3 cells cultured in 3D-C with or without Asn (0.1 mM), measured by quantitative RT-PCR and normalized to the non-treated condition. Student’s t test. **(L)** Confocal fluorescence microscopy images showing CD44 membrane localization in PC3 cells grown in standard 2D conditions, 3D-C, and 3D-C supplemented with Asn (0,1 mM). The image is representative of three independent experiments. **(M, N)** Western blot analysis of CD44 expression in PC3 cells grown in 3D-C and silenced for NARS1 (M) or GFPT1 (N), cultured with or without Asn (0.1 mM). Vinculin was used as a loading control. The images are representative of three independent experiments. **(O)** Total spheroid area of 3D-C PC3 cells silenced for the CD44 with or without Asn (0.1 mM). One-way ANOVA with Dunnett’s correction. **(P)** Hyaluronic acid adhesion assay on CD44-silenced PC3 cells grown in 3D-C with or without Asn (0.1 mM). One-way ANOVA with Dunnett’s correction. ns, not significant; *P < 0.05; **P < 0.01; ***P < 0.001; ****P < 0.0001. Data represent mean ± s.e.m. from at least three independent experiments.

Notably, CD44 protein levels increased in response to Asn in a dose-dependent manner (Fig. 6I), whereas depletion of extracellular Asn with ASNase abolished the Asn-induced upregulation in CD44 protein content (Fig. 6J). Loss of N-glycosylation has been shown to alter CD44 subcellular localization and promote its lysosomal degradation, ultimately compromising its function^44^. In keeping, CD44 mRNA levels did not change upon Asn supplementation in 3D-C (Fig. 6K), despite a marked increase in the relative protein levels. These data suggest that Asn supports increased CD44 translation and membrane stability via post-translational mechanisms, likely N-linked glycosylation, rather than a transcriptional regulation. In agreement, Asn stimulation specifically promoted CD44 localization to the extracellular membrane in 3D-C (Fig. 6L). In turn, preventing Asn incorporation into nascent proteins by silencing NARS1, or inhibiting protein N-glycosylation through GFPT1 silencing, reduced CD44 protein levels and abolished the Asn-mediated increase in CD44 expression (Fig. 6M, N), confirming the critical role of N-linked glycosylation in maintaining CD44 stability. Finally, CD44 silencing abrogated the Asn-mediated enhancement of both cell-cell clustering (Fig. 6O) and the adhesion to hyaluronic acid (Fig. 6P), underscoring its central role in mediating the pro-metastatic effects of Asn in bone and lung.

Collectively, these findings identify CD44 as a pivotal effector linking Asn-mediated protein N-glycosylation to the cell adhesive properties required for PC colonization of bone and lung ECM, highlighting its potential as a therapeutic target to prevent metastasis.

## DISCUSSION

Metastatic dissemination is a multistep process that requires cancer cells to dynamically adapt to the diverse and often hostile environments encountered along the metastatic cascade^45^. During this process, matrix detachment is accompanied by extensive metabolic remodeling^46–48^, which promotes resistance to anoikis and ferroptosis^49^ and supports the acquisition of stem-like traits that facilitate metastatic colonization^50^. Here, we identify a previously undescribed adaptive response of detached cells consisting in a metabolic shift toward the HBP, predisposing cells to increase protein glycosylation. Notably, alteration in metabolic pathways impacting in HBP activity and protein glycosylation have been already described to facilitate cancer metastases^51^. Our data further indicate that this adaptation becomes particularly advantageous upon arrival in Asn-rich environments, such as bone and lung, where increased Asn availability enables enhanced translation of specific subset of N-linked glycoproteins functional for successful colonization.

Once arrived at distant sites, metastatic cells are known to undergo further metabolic adaptations to cope with altered nutrient abundance and bioenergetic challenges^52^. However, studies addressing these adaptations have predominantly focused on metabolites as sources of energy or biosynthetic building blocks^53–55^. By contrast, considerably less attention has been given to the non-canonical roles of metabolites. Beyond supporting biomass and energy production, metabolites can function as intra- or extracellular signaling molecules^8,11^, act as epigenetic regulators^17^, or serve as substrates for post-translational modifications in metastatic cells^51^. Although Asn has been reported as a key modulator of metabolic homeostasis in cancer cells^15,23,56^ no metabolic adaptation accounted for its pro-metastatic effects in our model. Instead, Asn activates mTORC1 signaling and enhances protein synthesis in cells growing under anchorage-independent conditions, resulting in increased spheroid formation, independently of proliferation. Together with changes in the richness of specific tRNAs, which have been implicated in proteomic shifts toward a pro-metastatic state^57^, alterations in amino acid abundance influence translation efficiency by modulating their selective incorporation into nascent polypeptides, thereby shaping proteome architecture^58^. For instance, amino acid limitation has been shown to selectively impair tRNA^Gln^ charging, leading to reduced synthesis of core transcription factors whose protein sequences contain polyglutamine tracts^59^. Within this context, amino acid availability emerges as a key determinant in shaping proteome architecture^58^. Consistent with this concept, our data demonstrate that Asn does not indiscriminately enhance translation in metastatic cells but instead selectively sustains the synthesis of Asn-rich proteins. Accordingly, disrupting Asn-specific tRNA charging by silencing NARS1 abolished the Asn-dependent translational advantage under anchorage-independent conditions. In line with these findings, treatment of PC cells with ASNase has been shown to induce ribosomal pausing at Asn codons, highlighting a specific vulnerability of cancer cells to Asn availability during protein translation^30^. Reflecting this concept, ASNase treatment in our model consistently abolished the Asn-dependent enhancement of spheroid formation. In breast cancer, Asn availability has been shown to regulate metastatic progression through control of epithelial–mesenchymal transition (EMT)^60^. Proteomic analyses from that study further revealed that EMT-driving proteins are enriched in Asn residues, and that restricting Asn bioavailability impairs the expression of EMT-upregulated proteins at both translational and transcriptional levels. Although the mechanisms linking Asn availability to transcriptional regulation of EMT-associated genes remain unclear, our data provide a plausible explanation for the reported translational effects. Specifically, we find that elevated Asn content across proteins correlates with increased representation of consensus motifs for N-glycosylation. Because N-glycosylation is critical for the stability and function of multiple glycoproteins involved in EMT^61^, this relationship may help explain how Asn availability enhances the expression of EMT-associated proteins in breast cancer. Indeed, in our model, we found that Asn-rich proteins upregulated in bone and lung metastases are significantly enriched in predicted N-glycosylation sites. Exposure of cells grown under anchorage-independent conditions to elevated Asn levels resulted in a marked increase in global N-glycosylation. Conversely, genetic inhibition or tunicamycin-mediated blockade of N-glycosylation process completely abolished the Asn-mediated advantage in spheroid formation, establishing N-glycosylation as a functional downstream effector. From a functional perspective, we demonstrated that Asn-dependent enhanced protein N-glycosylation is essential for efficient cell-cell and cell-ECM adhesion, processes critical for metastatic seeding and colonization. Specifically, this transcriptional adaptation increases the aggregation capacity of metastatic spheroids and strengthens adhesion to bone- and lung-derived extracellular matrices. Importantly, prolonged Asn exposure during spheroid formation, rather than acute supplementation during adhesion assays, was required to confer these adhesive advantages, further supporting the notion that Asn drives durable translational remodeling rather than transient signaling events.

Notably, CD44, a heavily N-glycosylated HA receptor^62^, emerged as a key mediator of Asn-mediated metastatic colonization by promoting interactions with the bone and lung ECM. Asn exposure increases CD44 protein abundance and membrane localization without affecting CD44 mRNA levels, whereas blocking Asn incorporation into newly synthesized proteins or inhibiting N-glycosylation markedly reduced the Asn-dependent increase in CD44 protein. Consistent with our model, previous studies have shown that, in breast cancer, aggregation of DTC enhances intercellular CD44–CD44 homophilic interactions, thereby promoting metastatic outgrowth^63^. Moreover, CD44 plays a central role in maintaining cancer stemness through its interaction with HA and the activation of signaling pathways required for the acquisition and persistence of stem-like traits^64^. Enhanced CD44 exposure at the cell surface within Asn-rich metastatic niches may therefore represent an additional mechanism by which elevated extracellular Asn supports early metastatic colonization, fostering DTC clustering and reinforcing the self-renewal capacity of metastatic “seeds.” In this context, CD44-targeted therapeutic strategies (including monoclonal antibodies and HA-conjugated drug delivery systems) could be proposed as promising approaches to interfere with Asn-supported metastatic colonization^65,66^. Notably, several CD44-targeting agents have already entered clinical evaluation (NCT01358903, ^67^), and our findings provide a strong mechanistic rationale for their potential application in metastatic PC.

Another important translational implication of our findings is the possibility of limiting metastatic progression by reducing environmental Asn, either through dietary intervention or pharmacological depletion^68^. In *in vivo* PC metastasis model, we observed that dietary Asn restriction impairs metastatic colonization, with a selective effect in bone and lung. Consistently, culturing non-adherent cells in the presence of ASNase abrogated the advantages conferred by Asn supplementation, supporting the potential repurposing of this FDA/EMA-approved drug to interfere with metastatic disease in PC. ASNase is currently a standard component of chemotherapy regimens for acute lymphoblastic leukemia (ALL) and, in some cases, lymphoblastic lymphoma^69^, but its clinical use in solid tumors remains limited^70,71^. These limitations have been largely attributed to the metabolic plasticity of solid tumors and the emergence of resistance mechanisms, including upregulation of ASNS^72^. Importantly, our data suggest that metastatic PC cells colonizing bone and lung may be particularly vulnerable to extracellular Asn depletion due to the concurrent downregulation of ASNS under matrix-detached conditions. This vulnerability points to a potential therapeutic window during early metastatic colonization. In this context, systemic Asn depletion could selectively impair metastatic outgrowth without affecting normal tissues with intact ASNS expression.

In conclusion, this study identifies a niche-specific metabolic dependency that can be therapeutically exploited to limit skeletal and lung metastases in PC and potentially in other bone- and lung-tropic tumors, including breast cancer. More broadly, it provides a conceptual framework for investigating how tissue-specific nutrient landscapes shape metastatic tropism across tumor types.

## METHODS AND MATERIALS

### Cell Models

Human prostate cancer cells (PC3, RRID: CVCL_0035; DU145, RRID: CVCL_0105) were obtained from ATCC. All cells were maintained in high glucose (4.5 g/L) Dulbecco’s modified Eagle’s medium (DMEM) (ECB7501L; Euroclone) supplemented with 10% fetal bovine serum (#ECB4004L; Euroclone), 2 mmol/L L-glutamine and 1% penicillin/streptomycin (final concentration of 50 U/mL and 50 μ/mL respectively), at 37°C in a humidified atmosphere containing 5% CO_2_.

Metastatic cells were obtained from PC3-Luc^+^ cells (PC-3-Luc2: CRL-1435-LUC2) and PC3-GFP^+^ cells (RRID: CVCL_YJ90) (Innoprot). Cells were routinely tested for Mycoplasma contamination with the MycoAlert Mycoplasma Detection kit (#LT07–318; Lonza). 3D cultures were performed as previously described^16^. Briefly, agar (Sigma-Aldrich) was dissolved at 1% in distilled water and sterilized before being mixed with prewarmed (37 °C) starvation DMEM at a 1:1 ratio. The agar mixture was then transferred into six-well plates (3 ml per well) or p100 plates (10 ml per plate) and allowed to solidify at room temperature. PC cells were subsequently seeded on top of the base agar at a density of 18,000 cells per well (six-well plates) or 700,000 cells per plate (p100 plates) in serum-free culture medium supplemented with 2% dialyzed FBS (dFBS) (Slide-A-Lyzer dialysis cassettes, 3.5 kDa cutoff; Thermo Scientific), with additional treatments when indicated. All analyses were performed after 5 days of incubation. The 3D area of cell clusters was quantified using ImageJ software (NIH, USA).

The following inhibitors were used in this study: 0.25 U/ml L-asparaginase (#HY-P1923, BioCompareDBA), 1 mM Mitomycin C (#M4287, Sigma-Aldrich), 0.01 mM Rapamycin (#553210, Sigma-Aldrich), 0.1 mM Ethylenediaminetetraacetic acid (EDTA) (#E6758, Sigma-Aldrich), 0.1 mM Cycloheximide (#C7698, Sigma-Aldrich), 5ng/µl Tunicamycin (#T7765, Sigma-Aldrich).

### Metastatic Mouse Models

All animal procedures were approved by the Italian Ministero della Salute (Project License No. 762/2018) and carried out in accordance with institutional and national ethical guidelines, as approved by the Institutional Animal Care and Use Committee of the University of Florence. For tumor models, mice were randomly assigned before cancer cell injection. All analysis was performed in a blinded manner. Sample size was determined by power analysis (B = 0.8, p < 0.05) based on preliminary data and in compliance with the 3Rs principle: Replacement, Reduction, and Refinement.

Male SCID mice (NOD.CB17-Prkdc^scid/NCrHsd, Envigo), 6–8 weeks of age, were housed at the Centro Stabulazione Animali da Laboratorio (CESAL, University of Florence, Italy) under sterile conditions, with ad libitum access to food (standard diet, Mucedola Srl) and water. Humane endpoints were defined as follows: a maximum tumor volume of 1.8 cm³, impaired mobility, labored respiration, surgical site infection, or body weight loss exceeding 10% of the initial value. Mice were monitored regularly, and any animal displaying one or more of these signs was humanely euthanized.

1×10^⁵^ and 1×10^⁶^ PC3-Luc⁺ or PC3-GFP⁺ cells were injected via intracardiac (i.c.) or caudal intra-arterial (c.a.) routes, respectively, into NOD-SCID mice. For i.c. injections, a 100 µL bolus containing 1×10^⁵^ PC3-Luc⁺ cells were inoculated into the left ventricle under high-frequency ultrasound guidance using the VevoF2-LAZR-X imaging system (VisualSonics Fujifilm). During the procedure, animals were anesthetized with isoflurane (5% in oxygen for induction and 2% for maintenance). Cells were injected using a tuberculin syringe equipped with a 30-gauge needle. For imaging, a high-frequency ultrasound transducer with a 57–25 MHz bandwidth (UHF57x, VisualSonics, FUJIFILM) was used. Metastatic progression was quantified weekly *in vivo* using the IVIS Lumina S5 imaging system (Perkin Elmer, Waltham, MA, USA) starting from 4 weeks after PC cell injection. Before each imaging session, mice received an intraperitoneal injection of D-luciferin (Promega) at 100 mg/kg, dissolved in PBS (15 mg/ml). Groups of five animals were anesthetized with 2% isoflurane and imaged simultaneously using the IVIS Lumina S5 imaging system (Perkin Elmer). Bioluminescence signals were analyzed using the Living Image software (Perkin Elmer), and identical regions of interest (ROIs) were defined around the bone of each mouse. Data were expressed as total photon flux (photons/second) within each animal or in selected ROIs.

For the asparagine dietary deprivation experiment, mice were fed with a control diet (CD) or asparagine-deprived diet (ADD) (50g/mouse/week) starting from 2 weeks before PC cells injection. The experimental diets applied were provided by Mucedola Srl and were formulated as previously described in^60^. Briefly, the control diet contains all the essential amino acids, including asparagine (6 g/Kg). The asparagine-free diet has the same formulation of the control diet but lacks asparagine, compensated for a proportional addition of the other amino acids to reach an equal amount of amino acid content.

For metastatic cell isolation, mice were sacrificed two months after injection, and bone, lung, and liver tissues were collected. Single-cell suspensions were prepared and subjected to fluorescence-activated cell sorting (FACS) using a BD FACSMelody Cell Sorter (Becton Dickinson). Sorted cells were either seeded onto culture plates to establish metastatic cell derivatives or collected (1.000-5.000 cells) for RNA extraction in Extraction Buffer (Arcturus™ PicoPure™ RNA Isolation Kit, Thermo Fisher Scientific). Cells (50.000 each sample) were obtained from a primary tumor derived from subcutaneous engraftment into the flanks of mice (1×10^6^ PC3-GFP⁺ cells) were processed in parallel and used as controls.

### Transient transfections

Cells were transfected with a pool of siRNA targeting human GFPT1 (EHU082891; Merck Millipore), STT-A (EHU092531; Merck Millipore) and STT-B (EHU051881; Merck Millipore), CD44 (EHU094691; Merck Millipore) using Lipofectamine RNAiMAX transfection reagent (Thermo Fisher Scientific #13778150) according to the manufacturer’s instructions.

### Migration Assay

Migration assays were performed using 8-μm-pore transwell inserts (Corning #3428). Cells (3.5 × 10⁴ per well) were seeded in the upper chamber in serum-free medium and allowed to migrate for 16 h toward medium containing 2% dialyzed FBS, with or without 0.1 mM asparagine. After incubation, membranes were air-dried and stained with Diff-Quick (BD Biosciences #726443). Migrated cells on the lower surface were quantified by counting six randomly selected fields per membrane.

### Short-time aggregation assay

Cells were cultured for 5 days under the indicated experimental conditions, either as 3D-C or as standard 2D adherent cultures. After 5 days, cells were collected and thoroughly dissociated to obtain a single-cell suspension. An equal number of cells for each condition was then replated onto non-adherent plates previously coated with agar at 1% in distilled water at a 1:1 ratio in starvation DMEM. Cells were incubated for 6 hours under standard culture conditions, after which images were acquired, and the area of the newly formed cell clusters was quantified.

### Adhesion Assays

Plates were coated with Bone dECM Gel, Lung dECM Gel, or Liver dECM Gel Hydrogel Kits (Merck Sigma #CC170, #CC175, #CC174, respectively) according to the manufacturer’s instructions, or with collagen type I (10 μg/ml; rat tail, BD Biosciences #354236) or hyaluronic acid (0.5 mg/ml; Merck Sigma #924474). After 5 days of the indicated treatments,cells were collected from 3D-C cultures and seeded onto the coated plates for 15 minutes. Adherent cells were fixed in methanol, and images of each well were acquired. Cell numbers were quantified using ImageJ (NIH).

### KI-67 proliferation analysis

After 5 days of the indicated treatments, 3D-C cells were collected by centrifugation at 3,000 rpm for 5 minutes, washed with PBS, and fixed by adding cold 70% ethanol dropwise while vortexing. Cells were incubated on ice for at least 30 minutes, washed three times with cold PBS containing 1% BSA, and then stained with APC-conjugated Ki-67 antibody (0.03 μg, Thermo Fisher Scientific #17-5698-82, clone SolA15) diluted in FACS buffer for 30 minutes. Data were acquired on a FACSCanto II flow cytometer (BD Biosciences), and cell cycle profiles were analyzed using FlowJo with the Watson model or by manual gating.

### Seahorse analysis

After 5 days of the indicated treatments, cells from 3D-C were collected by centrifugation at 3,000 rpm for 5 minutes and then plated at a density of 2.5 × 10⁴ cells per well in XFe96 cell culture plates (6-8 technical replicates per condition) in 80 μl of standard medium supplemented with 2% dialyzed FBS. Cells were allowed to adhere at 37°C for 4-6 hours before further analysis. 1h before the analysis, media were replaced with XF Base Medium adjusted PH at 7.4. Cells were incubated for 1h at 37°C in atmospheric CO2 conditions to pre-equilibrate cells. OCR and ECAR analysis were performed using Seahorse XF Cell Mito Stress Test (Agilent #103,015-100) according to manufacturer’s instructions. Mitochondrial drugs were utilized as follows: 0.8 μM of oligomycin, 1 μM of FCCP, 1 μM of rotenone, and 1 μM of antimycin A were injected three times subsequently at the times indicated. Results were normalized to protein content.

### Tissue collection for metabolite extraction

At the conclusion of *in vivo* experiments, mice were euthanized by cervical dislocation. Blood was promptly collected via cardiac puncture into heparin-coated tubes, and plasma was separated by centrifugation at 1,000-2,000 × g for 10 minutes at 4°C. Primary tumors, lungs, tibial bones, liver and brain were excised, rinsed in cold 0.9% NaCl, and snap-frozen in liquid nitrogen. Samples were stored at -80°C until further processing. For metabolite extraction, frozen tissues were weighed (20-50 mg) and mechanically homogenized.

### GC-MS metabolomics analysis

Cells in 3D-C were collected by centrifugation at 3,000 rpm for 5 minutes, and the resulting pellets were resuspended in ice-cold 0.9% NaCl. Cells were then lysed in 800 μL of cold 80% methanol in HPLC-grade water (containing 1 µg/ml norvaline and 1.25 µg/ml glutarate as internal standards). Metabolites from culture media were extracted by adding 80 μL of cold solution of methanol 80% in HPLC-grade water (containing 01 µg/ml norvaline and 1.25 µg/ml glutarate as internal standards) to 20 μl of media previously centrifuged for 10 min at 1.200 rpm. Metabolites from plasma were extracted by adding 100 µl of cold solution of methanol 100% (containing 1 µg/ml norvaline and 1.25 µg/ml glutarate as internal standards) to 100 µl of plasma. Metabolites from tissues were extracted by resuspending homogenized tissues in 400 µl of cold solution of methanol 80% in HPLC-grade water (containing 1 µg/ml norvaline and 1.25 µg/ml glutarate as internal standards) and 400 µl of chloroform and sonicating them three times (10 s each) in ice. Samples were vortexed for 10 min at 4 °C and then centrifuged at 4 °C and 14,000 rpm for 10 min. Samples were dried by using a vacuum concentrator (Labconco). Polar metabolites were derivatized with 10 µl of a solution 40 µg/ml methoxyamine (Merck Sigma #226904) in pyridine (Merck Sigma #270970) for 30 min at 60 °C. GC-MS runs were performed by using an Intuvo 9000 GC / 5977B MS System (Agilent Technologies) equipped with an HP-5MS capillary column (30 m x 0.25 mm x 0.25 µm). 1 µl of each sample was injected in split or splitless mode using an inlet liner temperature of 240°C. Then, 45 µl of N-(tert-butyldimethylsilyl)-N-methyl-trifluoroacetamide, with 1% tert-butyldimethylchlorosilane (Merck Sigma #375934) were added in each sample and incubated for 60 min at 37 °C. GC runs were performed with helium as carrier gas at 1 ml/min. The GC oven temperature ramp was from 70°C to 280°C. The temperature of 70°C was held for 2min. Then, the first temperature ramp was from 70°C to 140°C at 3°C/min. The second ramp was from 140°C to 150°C at 1°C/min. The third temperature ramp was from 150°C to 280°C at 3°C/min. The measurement of metabolites was performed under electron impact ionization at 70 eV using a SIM mode. The ion source and transfer line temperature were set to 230°C and 290°C, respectively. For the determination of relative metabolite abundances, the integrated signal of all ions for each metabolite fragment was normalized by the signal from norvaline and sample protein content isolated by resuspending the protein layer in 50 μL NaOH 200 mM, vertexing for 15 min at 96°C, and centrifuging at 4°C 14.000 rpm for 15 min. Protein abundance was measured by BCA assay.

### LC-MS metabolomics analysis

Metabolites from tissues were extracted by resuspending homogenized tissues in 400 µl of solution of methanol/acetonitrile/H2O (50:30:20). For isotopomer distribution assay, fresh media containing [U-^13^C] glucose was added 24h before metabolite extraction to cells growing either as 3D-C or as standard 2D adherent cultures. Specifically, labeled media was formulated starting from DMEM without glucose (Gibco #11966025) supplemented with 2% dialyzed FBS, 1% penicillin and streptomycin (EuroClone #ECB3001D), 1mM Sodium Pyruvate (Thermo Scientific #11360070), and D-glucose (Merck Sigma #G8644) or [U-^13^C]-glucose (Cambrige Isotope laboratories #CLM-1396-1) to reach a final concentration 17 mM. Metabolites were extracted by lysing cells in a cold (-20°C) solution of methanol/acetonitrile/H2O (50:30:20). Samples were shaken at 4°C for 10 min, then centrifuged for 15 min at 16,000 × g. The supernatant was then collected and analyzed by LC-MS. 10 µl of each sample was loaded into a Dionex UltiMate 3000 LC System (Thermo Scientific Bremen, Germany) equipped with a C-18 column (Acquity UPLC-HSS T3 1. 8 µm; 2.1 x 150 mm, Waters) coupled to a Q Exactive Orbitrap mass spectrometer (Thermo Scientific) operating in negative ion mode. A step gradient was carried out using solvent A (10 mM TBA and 15 mM acetic acid) and solvent B (100% methanol). The gradient started with 0% of solvent B and 100% solvent A and remained at 0% B until 2 min post injection. A linear gradient to 32.3% B was carried out until 7 min and increased to 36.3% until 14 min. Between 14 and 26 minutes the gradient increased to 90.9% of B and remained at 90.9% B for 4 minutes. At 30 min the gradient returned to 0% B. The chromatography was stopped at 40 min. The flow was kept constant at 0.25 mL/min and the column was placed at 40°C throughout the analysis. The MS operated in full scan mode (m/z range: [70.0000-1050.0000]) using a spray voltage of 4.80 kV, capillary temperature of 300°C, sheath gas at 40.0, auxiliary gas at 10.0. The AGC target was set at 3.0E+006 using a resolution of 70000, with a maximum IT fill time of 512 ms. Data collection was performed using the Xcalibur software (Thermo Scientific). The data analyses were performed by integrating the peak areas (El-Maven–Polly-Elucidata). For the determination of relative metabolite abundances, the integrated signal of all ions for each metabolite fragment was normalized by sample protein content as previously described.

### Protein extraction and Western blot analysis

Cells were lysed in lysis buffer (30 mM Tris-HCl, pH 7.5; 240 mM NaCl; 50 mM KCl; 1% Triton X-100,) diluted 1:2 in bi-distilled water and supplemented with protease (Merck Sigma #P8340) and phosphatase (Merck Sigma #P5726) inhibitors. Protein lysates were centrifuged at 14,000 rpm and 4°C for 10 min.

Extracted proteins were quantified using a bicinchoninic acid (BCA) assay (Merck Sigma #BCA1). Subsequently, 20–25 μg of total proteins were loaded on SDS-PAGE gels and transferred to PVDF membranes (BioRad #1704157). Membranes were incubated for 1h at room temperature in blocking buffer (5% non-fat dry milk (SantaCruz Biotechnology #sc-2325) in PBS-Tween 0.1%), and then incubated at 4°C over-night with primary antibodies against either ATF-4 (Cell Signaling Technology #11815), ASNS (Cell Signaling Technology #92479S), Phospho-mTOR (Ser2448) (Cell Signaling Technology #5536S), mTOR (Cell Signaling Technology #2983), Phospho-S6K1(Thr389) (Cell Signalig Technology #9205), S6K1 (Cell Signaling Technology #9202), CD44 (Cell Signaling Technology #3570), GFPT1 (ABClonal #A20873), STT3a (Santa Cruz Biotechology #sc-390227), Vinculin (Santa Cruz Biotechology #sc-59803), β-Actin (Santa Cruz Biotechology #sc-47778). Primary antibodies were diluted 1:1000 in PBS-Tween 0.1% containing 5% BSA (Merck Sigma #A7906)). After washing in PBS-Tween 0.1%, membranes were incubated for 1 hour at room temperature with horseradish peroxidase-conjugated secondary antibodies anti-mouse (Santa Cruz Biotechology #sc-2005) or anti-rabbit (Santa Cruz Biotechology #sc-2357) diluted 1:2,500 in PBS-Tween 0.1% containing 1% BSA. Signal was detected using Clarity Western ECL Substrate (BioRad #1705061), and images were captured with a ChemiDoc MP Imaging System (BioRad). Band intensities were quantified using BioRad analysis software, with Vinculin or β-Actin serving as loading controls. All Western blot images are representative of at least three independent experiments.

### Protein synthesis (SUnSET)

Puromycin incorporation into nascent polypeptides was used to quantify protein synthesis, as previously described (PMID: 19305406). For 2D cultures, cells were seeded in 6-cm dishes and grown to 70–80% confluence at the end of the assay. For 3D-C cultures, cells were grown on agar-coated plates with or without Asn supplementation at the indicated concentrations. Five days after seeding, cells were washed with PBS and starved for 1 hour in Hanks’ Balanced Salt Solution (HBSS). Cells were then reactivated in medium containing 2% dialyzed FBS for 30 minutes. Puromycin (10 μg/ml) was added during the final 15 minutes of reactivation. Cells were lysed, and protein extracts were prepared using standard lysis buffer. Puromycin incorporation was assessed by western blot using an anti-puromycin antibody (Merck Sigma #MABE343)

### Concanavalin A staining

Cells were lysed in lysis buffer (50 mM Tris-HCl pH 7.5, 150 mM NaCl, 100 mM NaF, 2 mM EGTA, 1% Triton X-100) supplemented with protease (Merck Sigma #P8340) and phosphatase (Merck Sigma #P5726) inhibitors. Protein lysates were centrifuged at 14,000 rpm and 4°C for 10 min.

Extracted proteins were quantified using a bicinchoninic acid (BCA) assay (Merck Sigma #BCA1). Subsequently, 20–25 μg of total proteins were loaded on SDS-PAGE gels and transferred to PVDF membranes (BioRad #1704157). Membranes were incubated for 2 minutes in PBS containing 2% (v/v) TWEEN 20 at 20°C, rinsed twice in PBS, and then incubated with 1 μg/ml of lectine-peroxidase (Concanavalin A from Canavalia ensiformis (Jack bean) peroxidase conjugate, Merck Sigma # L6397) in PBS containing 0,05% (v/v) TWEEN 20, with 1mM CaCl_2_, and 1 mM MgCl_2_ for 16h at 20 °C. Images were captured with a ChemiDoc MP Imaging System (BioRad). Band intensities were quantified using BioRad analysis software, with Vinculin or β-Actin serving as loading controls. All images are representative of at least three independent experiments

### Proteomics analysis

Cells were washed with PBS and lysed in lysis buffer (15 mM Tris pH 7.5, 25 mM KCl, 120 mM NaCl, 0.5% Triton X-100) supplemented with protease and phosphatase inhibitor cocktail (Halt Protease Inhibitor Cocktail/Halt Phosphatase Inhibitor Cocktail, Thermo Fisher Scientific Waltham, MA, USA). Cell lysate was sonicated at 4 °C for 10″ three times. Afterwards, it was centrifuged at 15,000×g for 30 min. The resulting supernatant was accurately transferred to a new tube. Protein concentration was measured by the Bradford method (Bio-Rad, Hercules, CA, USA). Proteins extracts were diluted with 8 M urea in 50 mM ammonium bicarbonate (NH4HCO3) pH 8.8; 10µg of each sample were reduced with 200 mM 1,4-dithiothreitol (DTT) (Sigma-Aldrich), in 50mM NH4HCO3, for 1 h at 56 °C, and then alkylated with 200 mM 2-iodoacetamide (IAA) (Sigma-Aldrich), in 50mM NH4HCO3, for 30 min at room temperature in darkness. Samples were diluted to reduce the urea concentration below 1.4 M and digested overnight at 37 °C with sequencing-grade trypsin (Sigma-Aldrich) using a 1:30 (w/w) enzyme-to-protein ratio. Digestion was stopped by the addition of 1% TFA. Whole samples were dried down using a vacuum evaporator and stored at -80°C^73^. Liquid Chromatography Tandem Mass Spectrometry (LC-MS/MS) was carried out using an Easy nLC-1000 chromatograpy system coupled to an Exploris 480 mass spectrometer (Thermo Scientific, Bremen, Germany). Peptides separation was performed using a 15 cm, 75 μm i.d. reversed-phase column, in-house packed with 3 μm C18 silica particles (Dr Maisch). The gradient was generated using mobile phase A (0.1% FA and 2% ACN) and mobile phase B (0.1% FA and 80% ACN). Peptides separation was done at a flow rate of 230 nL/min using the following gradient: from 4% B to 44% B in 40 min, from 44% B to 100% B in 8 min, and from 100% B to 0% B in 8 min; then the column was cleaned for 5 min with 100% B. The peptides were subjected to electrospray ionization (ESI) followed by MS/MS. The mass spectrometer operated in DDA mode using a top-10 method. In detail, the MS full scan was 375–1400 m/z with a resolution of 60,000, AGC target ‘’custom’’ and maximum injection time of 50 ms. The mass window for the isolation of the precursor was 1.6 m/z, with a resolution of 30,000, an AGC target “custom” and a maximum injection time of 120ms; HCD fragmentation was set at normalized collision energy of 30 and dynamic exclusion of 20 s. Raw files were processed by MaxQuant software version 2.1.4 using the Andromeda search engine against Human-refprot-isoforms.fasta in the NCBI. Database searches were performed using a precursor mass tolerance of 10 ppm and MS/MS tolerance of 0.02 Da. For peptide identification, two missed cleavages were allowed. The search included carbamidomethylation of cysteine as a fixed modification and oxidation of methionine as a variable modification. Potential common contaminants, proteins identified in the reverse database, and proteins identified only by site were excluded. Protein quantification was based on the MaxQuant label-free algorithm using unique and razor peptides and the matching-between-run feature selection (LFQ)^74^. Statistical analysis was performed using Perseus software version 2.0.11. Data were log2 transformed based on LFQ intensity, considering a minimum of 70% valid values in at least one group. For statistical analysis and the identification of significant proteins, a false discovery rate (FDR) of 0.05% and a fold change (FC) of 2 were established^75^. Protein-protein interaction networks and functional enrichment analyses were performed using IPA Ingenuity software, considering significant pathways with a p-value <0.01%.

### Protein annotation and N-glycosylation analysis

Proteins identified by proteomics analysis or proteins encoded by genes identified through RNA-seq analysis were annotated and analyzed as follows. For data derived from RNA-seq analysis, gene symbols derived from the significative differential genes between bone metastases/lung metastases/liver metastases vs primary tumor comparison (p-value threshold of less than 0.05) were converted into protein identifiers using the UniProt ID mapping tool (https://www.uniprot.org/id-mapping). Gene names were mapped to UniProtKB entries restricted to Homo sapiens. Only reviewed entries (Swiss-Prot) were retained to ensure high-confidence protein annotation. Mapped results were downloaded, including the following fields: protein length, amino acid sequence, glycosylation annotations, and post-translational modification information. For each protein sequence, protein lengths were obtained directly from the UniProt annotations and the relative abundance of Asn was calculated by counting the number of N residues within each protein sequence and normalizing this value to the total amino acid content of the corresponding protein.

N-glycosylation sites were analyzed by identifying N-linked glycosylation motifs (N-X-S/T, where X ≠ P). Lists of predicted N-glycosylation sites were generated separately for upregulated and downregulated proteins. The number of N-linked glycosylation sites per protein (“N-linked count”) was calculated automatically using custom scripts in RStudio. Glycosylation counts were subsequently normalized to protein length to account for differences in sequence size.

### RNA extraction and quantitative real-time PCR (qRT-PCR) analysis

Total RNA was extracted using the RNAeasy Plus Mini Kit (Qiagen, #74134) and quantified by NanoDrop One (Thermo Fisher Scientific). cDNA was synthesized from 1000 ng of total extracted RNA using the iScript cDNA Synthesis Kit (BioRad, #1725035) and the T100 ThermalCycler (BioRad), according to manufacturer’s instructions. qRT-PCR was performed to quantify mRNA levels of selected targets by CFX96 Touch Real-Time PCR Detection System (BioRad) using the TaqMan Universal PCR Master Mix (Applied Biosystem, #4440040), according to manufacturer’s instructions. The following probes were used: OST4 (Thermo Fisher Scientific #Hs00951309) GFTP1 (Thermo Fisher Scientific #Hs00899865), CD44 (Thermo Fisher Scientific #3570). Data are reported as relative quantity with respect to reference samples determined with the 2-ΔΔCt method by CFX Maestro software (BioRad). Data were normalized on TATA-Box Binding Protein (TBP, #Hs00427620, Thermo Fisher Scientific).

### RNA sequencing analysis

Total RNA sequencing (RNA-seq) was performed to characterize the transcriptional profiles of metastatic tumor cells isolated *in vivo* from primary tumors (PT) and from bone, lung, and liver metastatic sites. Total RNA was extracted using the PicoPure RNA Isolation Kit (ThermoFisher, #KIT0204) following the manufacturer’s instructions, which is optimized for low-input samples. Library preparation was carried out by IGA Technology Services (Italy) using an ultra-low input RNA-seq workflow. cDNA synthesis and library construction were performed using Universal Plus mRNA-Seq kit (Tecan Genomics, Redwood City, CA) according to the provider’s guidelines. Sequencing libraries were quantified, pooled, and sequenced on NovaSeq 6000 platform (Illumina, San Diego, CA) generating paired-end 150 bp reads, with a sequencing depth of approximately 60 million read pairs per sample. Raw sequencing data underwent quality control with FastQC, and adapters and low-quality bases were removed. RNA-seq data were processed using the Transcriptomestic pipeline (v1.0; available at github.com/littleisland8/Transcriptomestic), a Snakemake-based workflow designed for reproducible transcriptomic analysis. The quality of raw sequencing reads was assessed using FastQC (v0.12.1) (questa citazione non ha PMID, Andrews, S. (2010) FastQC: A Quality Control Tool for High Throughput Sequence Data. https://www.bioinformatics.babraham.ac.uk/projects/fastqc/). Prior to alignment, adapter sequences and low-quality bases were removed using fastp^76^. High-quality reads were then mapped to the human reference genome (available through the 1000 Genomes FTP site ftp://ftp.1000genomes.ebi.ac.uk/vol1/ftp/technical/reference/GRCh38_reference_genome/GRCh38_full_analysis_set_plus_decoy_hla.fa) using STAR (v2.7.11a;^77^) with default parameters. Mapped reads were quantified at the gene level using HTSeq-count (v2.0.2;^78^), using the Ensembl GRCh38 gene annotation (v111). Differential gene expression analysis was conducted using the DESeq2 R package (v1.46.0;^79^). Pathway enrichment analysis was performed using the DOSE R package (v4.0.1;^80^). Gene sets were filtered to include only those containing between 2 and 500 genes. For all analyses, statistical significance was defined by a p-value and q-value (False Discovery Rate, FDR) threshold of less than 0.05

### Confocal Imaging

After 5 days of the indicated treatments, cells from 3D-C cultures were collected and seeded onto glass coverslips previously coated with collagen type I (10 μg/ml; rat tail, BD Biosciences #354236). Cells were fixed with ice-cold methanol for 5 minutes, washed twice with PBST (0.1% Tween-20), and blocked in IFF buffer (1% BSA, 2% FCS in PBS). Cells were permeabilized with 0.25% Triton X-100 in PBS for 5 minutes, washed twice with PBS, and incubated overnight at 4°C with primary antibody against CD44 (Cell Signaling Technology #3570) diluted 1:100 in IFF. After three washes with PBST cells were incubated with Alexa Fluor 488-conjugated secondary antibody (Thermo Fisher Scientific #A-11001, rabbit) for 1 hour at room temperature. Nuclei were stained with DAPI (Thermo Fisher Scientific #D3571) for 10 minutes at room temperature. Three to five representative images per sample were acquired using a Leica TCS SP8 confocal microscope with LAS-AF software.

### Immunohistochemistry (IHC) analysis

Individual specimens from human primary tumors and bone metastasis were dissected and fixed in 4% PFA for histology and IHC analysis. PCa patients-derived FFPE were provided by Anatomical Pathology Unit from University Hospital Careggi (Florence, Italy). The Leica BOND-RXm automated system (Leica Biosystems) was used for the staining of 5 m-sections with CD44 antibody (1:100; Cell Signaling Technology #3570), upon their specificity testing on negative and positive tissue samples. Slides were developed with 3’,3-diaminobenzidine (DAB) (Leica Biosystems) and counterstained with hematoxylin. Images were acquired by using a slide scanner (AperioLV1; Leica Biosystems) and analyzed with ImageScope software (RRID:SCR_020993).

### Immunofluorescence analysis

Lung metastases were identified by pan-cytokeratin immunofluorescence staining. FFPE lung sections were processed using the Leica BOND-MAX™ automated staining system (Leica Microsystems), followed by standard washing steps and incubation with the primary antibody (pan-cytokeratin, Abcam, #ab234297). Sections were then incubated with Alexa Fluor 488–conjugated rabbit secondary antibody (Thermo Fisher Scientific, #A-11008) for 1 h at room temperature. Images (three to five fields per sample) were acquired using a TCS SP8 confocal microscope (Leica Microsystems) equipped with LAS-AF acquisition software, and metastatic cells were quantified per field.

### Statistical analysis

Data analysis was performed using Prism 10.4 (GraphPad software). All experiments presented were repeated in at least three independent biological replicates, with graphs showing circled-shaped points indicating technical and/or biological replicates. Data are presented as mean ± s.e.m. Unless stated otherwise, comparisons between 2 groups were made using the two-tailed, unpaired Student’s t-test. Comparisons between multiple groups were made using one-way or two-way analysis of variance (ANOVA). Tukey, or Šidák post-testing analyses with a confidence interval of 95% were used for individual comparisons as reported in figure legends. P values <0.05 were considered statistically significant: **** <0.0001, ***<0.001, **<0.01, *<0.05, ns: not statistically significant. When the comparison was not biologically relevant, no indication was reported in the figures.

## ACKNOWLEDGEMENTS

The work was funded by *Fondazione Associazione Italiana Ricerca sul Cancro (AIRC) ETS* and *Fondazione Cassa di Risparmio di Firenze* (grant Multiuser 19515 to PC, grant IG 24731 to PC). The data presented in the current study were in part generated using grants by European Union, National Recovery and Resilience Plan, Mission 4 Component 2—Investment 1.4—National Center for Gene Therapy and Drugs based on RNA Technology—NextGenerationEU—Project Code CN00000041-CUP B93D21010860004 (to EG) and Creation and strengthening of “innovation ecosystems”, construction of “territorial R&D leaders”—TUSCANY HEALTH ECOSYSTEM (THE) NextGenerationEU—Project Code ECS_00000017-CUP B83C22003920001 (to PC). The research was partially funded by the Italian Ministry of University and Research within the PRIN2022 PNRR and PRIN2022 programs (Progetti di Ricerca di Rilevante Interesse Nazionale 2022) in the framework of the National Recovery and Resilience Plan (PNRR), Mission 4 - Component 2 - Investment 1.1 “Research Projects of Relevant National Interest (PRIN), funded by the European Union – NextGenerationEU – Project Code P2022CE7SP - CUP B53D23033040001 and Project Code 2022F5JLSE - CUP B53D23021510006, respectively (to EG).

We thank Prof. Gabriella Nesi for providing human FFPE PCa bone metastasis samples. We thank the collaborators of the VIB Metabolomics Core Leuven for their valuable contribution to the research presented in this paper

## AUTHOR CONTRIBUTIONS

Conceptualization: EP, LI, EG, PC. Methodology: EP, LI, MI, GC, ML, SP, SR, AS. Investigation: EP, LI, MI, GB, GV. Visualization: EP, LI, EG, PC. Supervision: EG, PC. Writing-original draft: EP.

## DATA AVAILABILITY

The datasets generated and/or analyzed during the current study are available in the Gene Expression Omnibus (GEO) repository under accession number GSE320461.

---

**Suppl. Fig. 1.**
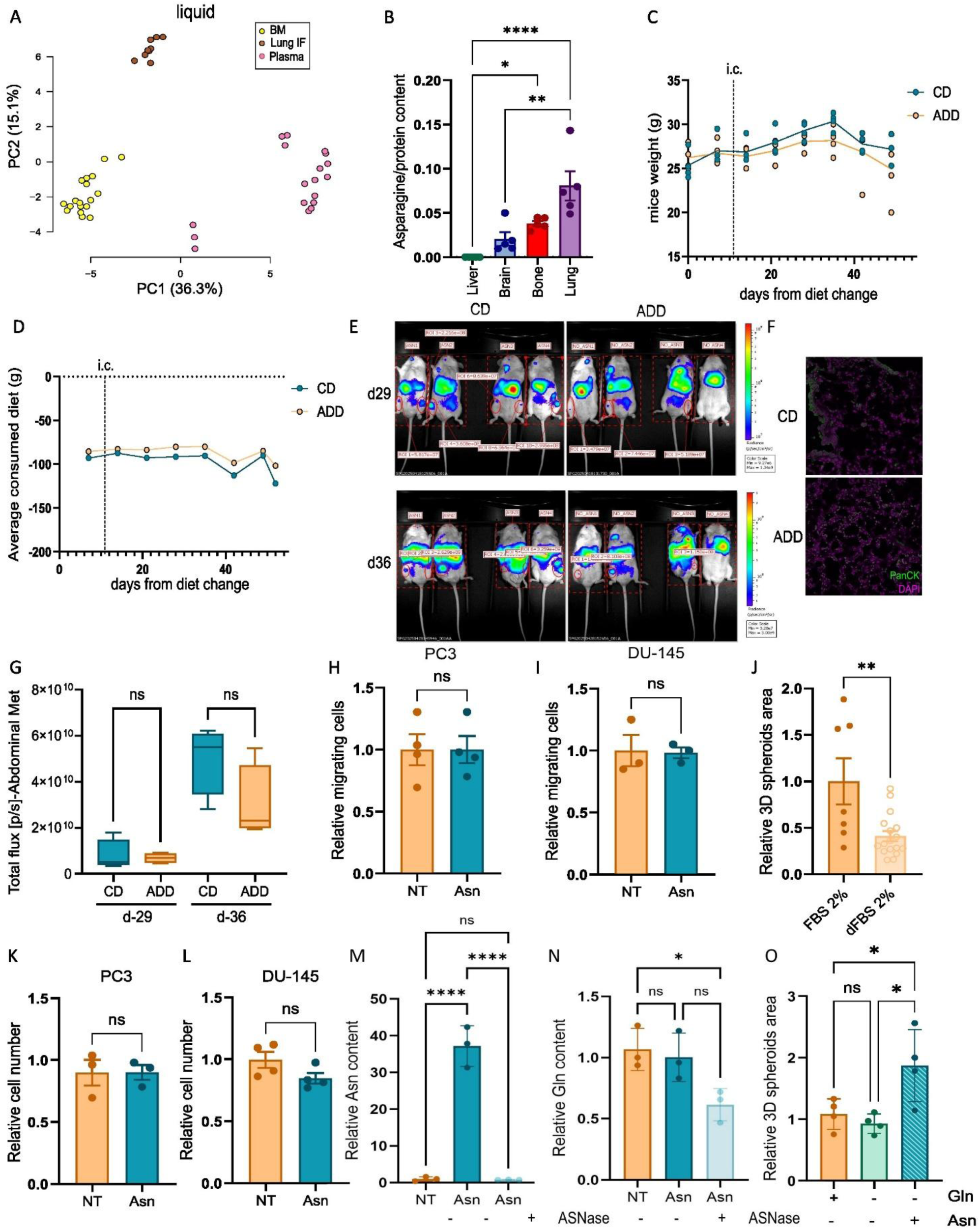
**(A)** Principal component analysis (PCA) of global metabolomic profiles from plasma, lung interstitial fluid (IF), and bone marrow collected from healthy NOD-SCID mice (n = 6) and analyzed by LC-MS. The PCA scatter plot shows PC1 (x-axis), accounting for 36.3% of the total variance, and PC2 (y-axis), explaining 15.1% of the variance. **(B)** Asn levels across tissues in healthy NOD-SCID mice (n = 5), quantified by GC–MS and normalized to total protein content. One-way ANOVA with Tukey’s correction. **(C)** Body weight changes in NOD-SCID mice fed an Asn-deprived diet (ADD) or control diet (CD). **(D)** Average daily food intake (g) of NOD-SCID mice receiving ADD or CD. i.c. denotes the day of intracardiac injection. **(E)** IVIS images of metastases formed after 29 or 36 post i.c. injection of Luc+ PC3 cells in NOD-SCID mice. ROIs corresponding to bone-specific metastasis are indicated by red circles and total photon flux (photons/second) within selected ROIs is indicated. **(F)** Representative immunofluorescence images for human Pancytokeratin (green) assessed in lung tissues derived from NOD-SCID mice receiving ADD or CD. Nuclei were stained with DAPI (pseudocolor magenta). **(G)** Quantification of abdominal metastases at days 29 and 36 after cancer cell injection, expressed as bioluminescent signal (photons/s). Abdominal metastases were measured within abdominal-specific ROIs. One-way ANOVA with Sidak’s correction. **(H-I)** Relative migration of PC3 (H) and DU-145 (I) cells assessed using Boyden chamber assays under nutrient-restricted conditions (2% dFBS) with or without Asn supplementation. Student’s t-test. **(J)** Total spheroid area of PC3 3D-C maintained in 2% FBS, dialyzed or not. Student’s t-test. **(K-L)** Relative cell number of PC3 (K) and DU-145 (L) cells cultured under standard 2D conditions for 5 days in the presence or absence of Asn (0.1 mM). Student’s t-test. **(M)** Relative Asn levels in DMEM supplemented or not with Asn (0.1 mM) and incubated for 5 days with or without L-asparaginase (ASNase, 0.25 U/ml). One-way ANOVA with Sidak’s correction. **(N)** Relative Gln levels in DMEM supplemented or not with Asn (0.1 mM) and incubated for 5 days with or without L-asparaginase (ASNase, 0.25 U/ml). One-way ANOVA with Sidak’s correction. **(O)** Total spheroid area of PC3 3D cultures grown with or without Asn (0.1 mM) in the presence or not of glutamine (Gln, 2 mM). One-way ANOVA with Sidak’s correction. ns, not significant; *P < 0.05; **P < 0.01; ***P < 0.001; ****P < 0.0001. Data represent mean ± s.e.m. from at least three independent experiments.

**Suppl. Fig. 2.**
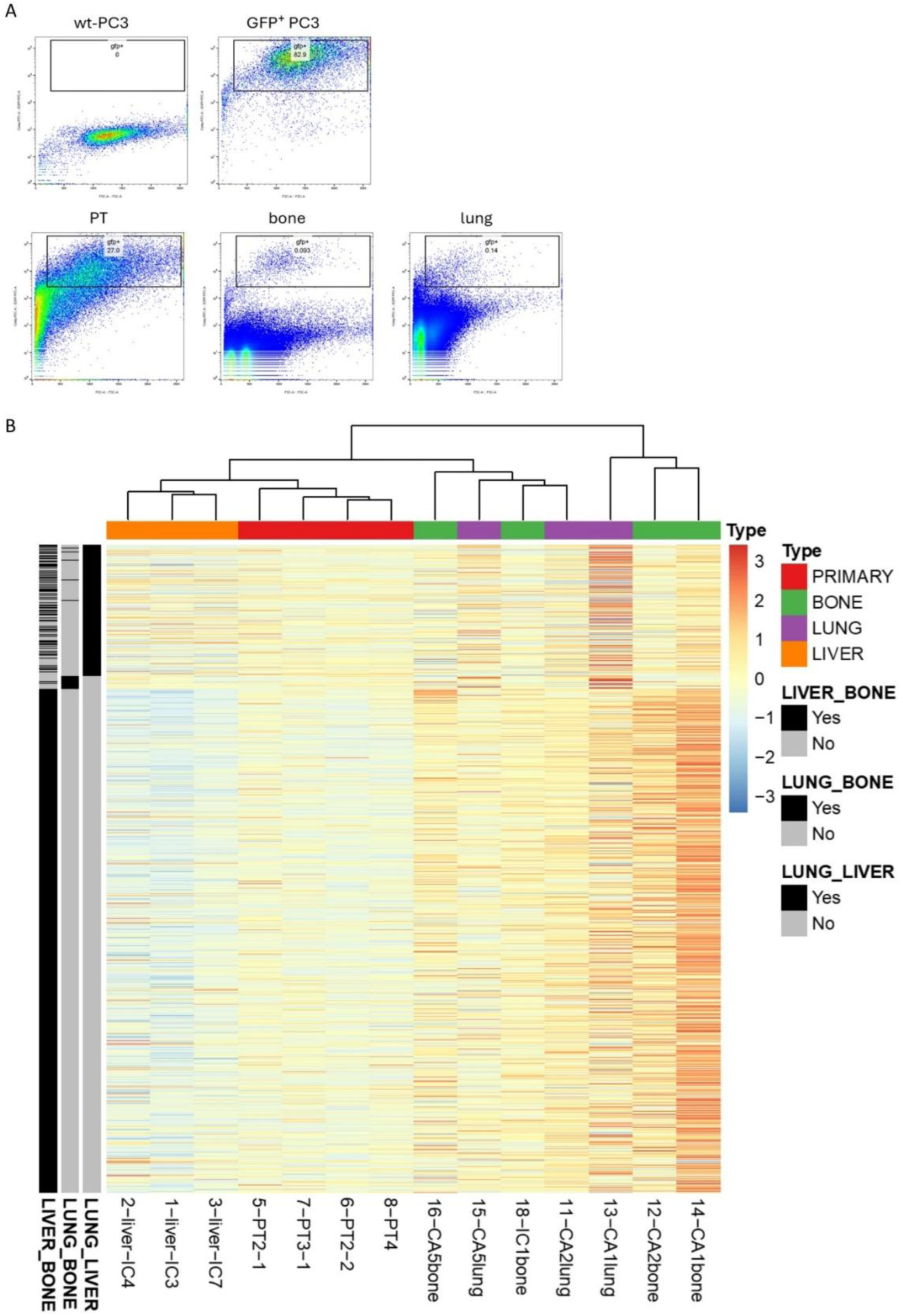
**(A)** Representative images of FACS-sorting selection of GFP⁺ cells isolated from primary tumors, bone, and lungs of NOD-SCID mice. Wild-type NOD-SCID mice injected i.c. or via the caudal artery (c.a.) with GFP⁺ PC3 cells were used as negative controls. **(B)** Heatmap and hierarchical clustering of RNA-seq gene expression profiles from metastatic PC3 cells obtained as described in Fig. 2M, together with cells from the corresponding primary tumor (PT). The heatmap displays normalized gene expression levels across samples, with rows representing genes and columns representing individual samples. Unsupervised hierarchical clustering reveals organ-specific transcriptional signatures. Color annotations indicate sample type (PT, bone, lung, or liver metastases). Side bars denote gene sets differentially associated with liver-bone, lung-bone, and lung-liver comparisons.

**Suppl. Fig. 3.**
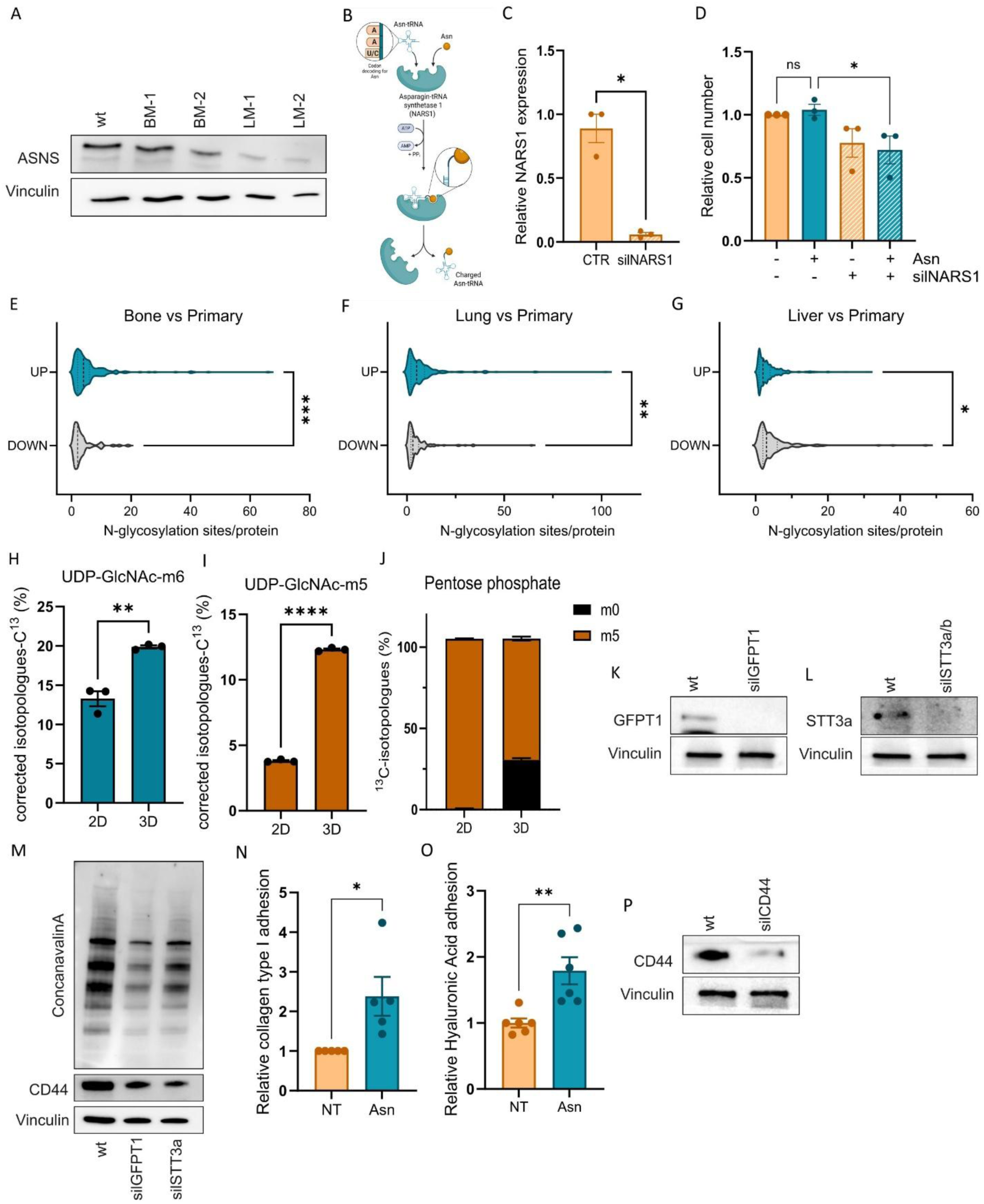
**(A)** Western blot analysis of ASNS in metastatic cells isolated from bone (B-M-1, B-M-2) and lung (L-M-1, L-M-2) lesions. Vinculin was used as loading control. The image is representative of three independent experiments. **(B)** Schematic representation of asparaginyl-tRNA synthetase 1 (NARS1) mechanism of action. **(C)** NARS1 mRNA levels in PC3 cells following NARS1 silencing. Cells were transfected with NARS1-targeting small interfering RNA (siRNA) or negative control, and mRNA levels were evaluated after 5 days of incubation in 3D cultures (3D-C) by quantitative RT-PCR, using scramble-transfected cells as reference. **(D)** Relative cell number of PC3 cells silenced for NARS1 and cultured under standard 2D conditions for 5 days in the presence or absence of Asn (0.1 mM). One-way ANOVA with Sidak’s correction. **(E-G)** Number of putative N-glycosylation sites in proteins encoded by genes up- or down-regulated in metastatic cells derived from bone (E), lung (F), and liver (G) relative to primary tumor (PT). RNA-seq analysis was conducted as described in Fig. 2M. Values are expressed relative to total protein number. **(H-I)** Fractional enrichment of UDP-GlcNAc isotopologues. PC3 cells were grown in 2D or 3D-C for 5 days and subsequently incubated in a medium containing U-^13^C-glucose for 24h. Labeling enrichment was evaluated by LC-MS analysis, and isotopologue abundance is reported as relative to total UDP-GlcNAc amount. Welch’s *t* test. **(J)** Labeling (m+5) enrichment of penotose phosphates from U-^13^C-glucose in PC3 cells growing in 2D or 3D-C for 5 days and subsequently incubated in a medium containing U-^13^C-glucose for 24h. Labeling enrichment was evaluated by LC-MS analysis. **(K, L, P)** Western blot analysis of GFPT1 (K), STT3a (L), and CD44 (P) expression in PC3 cells silenced for GFPT1, STT3a/b, and CD44 respectively after 48h of gene silencing. Vinculin was used as a loading control. The image is representative of three independent experiments. **(M)** Concanavalin A lectin binding assay performed on lysates from PC3 cells silenced or not for GFPT1 or STT3a/b and cultured in 3D-C for 5 days. Immunoblot for vinculin was used to confirm equal protein loading across samples. The image is representative of three independent experiments. **(N-O)** Adhesion of PC3 3D-C to collagen type I (L) and hyaluronic acid (M). Cells were cultured with Asn (0,1 mM) for 5 days and allowed to adhere for 15 min to plates coated with matrix components as reported. Adherent cells were quantified and data are shown relative to untreated cells. Welch’s t-test.

**Suppl. Tab. 1 3D-proteins** Perseus output results of proteins identified in PC3 cells grown in 3D-C in the absence or presence of Asn (0.1 mM). Sheet1: The table details for each identified protein: protein id, protein name, gene name, number of peptides, number of razor unique peptides, number of unique peptides, sequence coverage [%], razor peptides sequence coverage [%], unique sequence coverage [%], mol. weight [kDa], log2 of LFQ intensity for each replicate of analyzed sample, Significant proteins (FDR<0,05) are marked with*, Difference of treated cells vs untreated and -Log10(P-value) of the difference. Sheet 2: The table lists proteins with significantly different amounts (DEPs) between experimental conditions FDR<0.05 and fold-change cutoffs of 2.

**Suppl. Tab. 2 2D-proteins** Perseus output results of proteins identified in PC3 cells grown in 2D in the absence or presence of Asn (0.1 mM). Sheet1: The table details for each identified protein: protein id, protein name, gene name, number of peptides, number of razor unique peptides, number of unique peptides, sequence coverage [%], razor peptides sequence coverage [%], unique sequence coverage [%], mol. weight [kDa], log2 of LFQ intensity for each replicate of analyzed sample, Significant proteins (FDR<0,05) are marked with*, Difference of treated cells vs untreated and -Log10(P-value) of the difference. Sheet2: The table lists proteins with significantly different amounts (DEPs) between experimental conditions FDR<0.05 and fold-change cutoffs of 2.

**Supp. Tab. 3 3D exclusive proteins** Perseus output results of proteins identified exclusively in PC3 cells grown in 3D in the absence or presence of Asn (0.1 mM). The table lists proteins with significantly different amounts (DEPs) between experimental conditions FDR<0.05 and fold-change cutoffs of 2. The table details for each identified protein: protein id, protein name, gene name, number of peptides, number of razor unique peptides, number of unique peptides, sequence coverage [%], razor peptides sequence coverage [%], unique sequence coverage [%], mol. weight [kDa], log2 of LFQ intensity for each replicate of analyzed sample, Significant proteins (FDR<0,05) are marked with*, Difference of treated cells vs untreated and -Log10(P-value) of the difference.

## BIBLIOGRAPHY

1. Wong, S. K. et al. Prostate Cancer and Bone Metastases: The Underlying Mechanisms. Int. J. Mol. Sci. 20, (2019).

2. Bubendorf, L. et al. Metastatic patterns of prostate cancer: an autopsy study of 1,589 patients. Hum. Pathol. 31, 578–83 (2000).

3. Gao, Y. et al. Metastasis Organotropism: Redefining the Congenial Soil. Dev. Cell 49, 375–391 (2019).

4. Pranzini, E., Ippolito, L., Pardella, E., Giannoni, E. & Chiarugi, P. Adapt and shape: metabolic features within the metastatic niche. Trends Endocrinol. Metab. 36, 205–218 (2025).

5. Ngo, B. et al. Limited Environmental Serine and Glycine Confer Brain Metastasis Sensitivity to PHGDH Inhibition. Cancer Discov. 10, 1352–1373 (2020).

6. Christen, S. et al. Breast Cancer-Derived Lung Metastases Show Increased Pyruvate Carboxylase-Dependent Anaplerosis. Cell Rep. 17, 837–848 (2016).

7. Shinde, A., Wilmanski, T., Chen, H., Teegarden, D. & Wendt, M. K. Pyruvate carboxylase supports the pulmonary tropism of metastatic breast cancer. Breast Cancer Res. 20, 76 (2018).

8. Rinaldi, G. et al. In Vivo Evidence for Serine Biosynthesis-Defined Sensitivity of Lung Metastasis, but Not of Primary Breast Tumors, to mTORC1 Inhibition. Mol. Cell 81, 386–397.e7 (2021).

9. Elia, I. et al. Breast cancer cells rely on environmental pyruvate to shape the metastatic niche. Nature 568, 117–121 (2019).

10. Karno, B., Edwards, D. N. & Chen, J. Metabolic control of cancer metastasis: role of amino acids at secondary organ sites. Oncogene 42, 3447–3456 (2023).

11. Doglioni, G. et al. Aspartate signalling drives lung metastasis via alternative translation. Nature 638, 244–250 (2025).

12. Todd, V. M. & Johnson, R. W. Hypoxia in bone metastasis and osteolysis. Cancer Lett. 489, 144–154 (2020).

13. Edwards, D. N. Amino Acid Metabolism in Bone Metastatic Disease. Curr. Osteoporos. Rep. 21, 344–353 (2023).

14. Zhang, W. et al. The bone microenvironment invigorates metastatic seeds for further dissemination. Cell 184, 2471–2486.e20 (2021).

15. Krall, A. S., Xu, S., Graeber, T. G., Braas, D. & Christofk, H. R. Asparagine promotes cancer cell proliferation through use as an amino acid exchange factor. Nat. Commun. 7, 11457 (2016).

16. Elia, I. et al. Proline metabolism supports metastasis formation and could be inhibited to selectively target metastasizing cancer cells. Nat. Commun. 8, 1–11 (2017).

17. Altea-Manzano, P. et al. A palmitate-rich metastatic niche enables metastasis growth via p65 acetylation resulting in pro-metastatic NF-κB signaling. *Nat*. Cancer 4, 344–364 (2023).

18. Pilley, S. E. et al. Loss of attachment promotes proline accumulation and excretion in cancer cells. Sci. Adv. 9, eadh2023 (2023).

19. Kuchimaru, T. et al. A reliable murine model of bone metastasis by injecting cancer cells through caudal arteries. Nat. Commun. 9, 2981 (2018).

20. Zhang, J. et al. Asparagine plays a critical role in regulating cellular adaptation to glutamine depletion. Mol. Cell 56, 205–218 (2014).

21. Krall, A. S. et al. Asparagine couples mitochondrial respiration to ATF4 activity and tumor growth. Cell Metab. 33, 1013–1026.e6 (2021).

22. Pavlova, N. N. et al. As Extracellular Glutamine Levels Decline, Asparagine Becomes an Essential Amino Acid. Cell Metab. 27, 428–438.e5 (2018).

23. Krall, A. S., Xu, S., Graeber, T. G., Braas, D. & Christofk, H. R. Asparagine promotes cancer cell proliferation through use as an amino acid exchange factor. Nat. Commun. 7, 11457 (2016).

24. Meng, D. et al. Glutamine and asparagine activate mTORC1 independently of Rag GTPases. J. Biol. Chem. 295, 2890–2899 (2020).

25. Zoncu, R., Efeyan, A. & Sabatini, D. M. mTOR: from growth signal integration to cancer, diabetes and ageing. Nat. Rev. Mol. Cell Biol. 12, 21–35 (2011).

26. Panwar, V. et al. Multifaceted role of mTOR (mammalian target of rapamycin) signaling pathway in human health and disease. Signal Transduct. Target. Ther. 8, 375 (2023).

27. Micalizzi, D. S., Ebright, R. Y., Haber, D. A. & Maheswaran, S. Translational Regulation of Cancer Metastasis. Cancer Res. 81, 517–524 (2021).

28. McCool, E. N. et al. Deep top-down proteomics revealed significant proteoform-level differences between metastatic and nonmetastatic colorectal cancer cells. Sci. Adv. 8, eabq6348 (2022).

29. Minn, A. J. et al. Genes that mediate breast cancer metastasis to lung. Nature 436, 518–24 (2005).

30. Loayza-Puch, F. et al. Tumour-specific proline vulnerability uncovered by differential ribosome codon reading. Nature 530, 490–4 (2016).

31. Sabbía, V. et al. Trends of amino acid usage in the proteins from the human genome. J. Biomol. Struct. Dyn. 25, 55–9 (2007).

32. He, M., Zhou, X. & Wang, X. Glycosylation: mechanisms, biological functions and clinical implications. Signal Transduct. Target. Ther. 9, 194 (2024).

33. Breitling, J. & Aebi, M. N-linked protein glycosylation in the endoplasmic reticulum. Cold Spring Harb. Perspect. Biol. 5, a013359 (2013).

34. Paneque, A., Fortus, H., Zheng, J., Werlen, G. & Jacinto, E. The Hexosamine Biosynthesis Pathway: Regulation and Function. Genes (Basel). 14, (2023).

35. Moseley, H. N. B., Lane, A. N., Belshoff, A. C., Higashi, R. M. & Fan, T. W. M. A novel deconvolution method for modeling UDP-N-acetyl-D-glucosamine biosynthetic pathways based on (13)C mass isotopologue profiles under non-steady-state conditions. BMC Biol. 9, 37 (2011).

36. Reyes-Oliveras, A., Ellis, A. E., Sheldon, R. D. & Haab, B. Metabolomics and 13C labelled glucose tracing to identify carbon incorporation into aberrant cell membrane glycans in cancer. *Commun*. Biol. 7, 1576 (2024).

37. Fraser, J. R., Laurent, T. C. & Laurent, U. B. Hyaluronan: its nature, distribution, functions and turnover. J. Intern. Med. 242, 27–33 (1997).

38. Kylmälä, T. et al. Type I collagen degradation product (ICTP) gives information about the nature of bone metastases and has prognostic value in prostate cancer. Br. J. Cancer 71, 1061–4 (1995).

39. Bayne, E. F. et al. High-Throughput Extracellular Matrix Proteomics of Human Lungs Enabled by Photocleavable Surfactant and diaPASEF. J. Proteome Res. 23, 2908–2918 (2024).

40. Sroga, G. E., Karim, L., Colón, W. & Vashishth, D. Biochemical characterization of major bone-matrix proteins using nanoscale-size bone samples and proteomics methodology. Mol. Cell. Proteomics 10, M110.006718 (2011).

41. Arteel, G. E. & Naba, A. The liver matrisome - looking beyond collagens. JHEP Rep. 2, 100115 (2020).

42. Arriazu, E. et al. Extracellular matrix and liver disease. Antioxid. Redox Signal. 21, 1078–97 (2014).

43. Ponta, H., Sherman, L. & Herrlich, P. A. CD44: from adhesion molecules to signalling regulators. Nat. Rev. Mol. Cell Biol. 4, 33–45 (2003).

44. Cheng, Q. et al. N-glycosylation at N57/100/110 affects CD44s localization, function and stability in hepatocellular carcinoma. Eur. J. Cell Biol. 102, 151360 (2023).

45. Senft, D. & Ronai, Z. E. A. Adaptive Stress Responses During Tumor Metastasis and Dormancy. Trends Cancer 2, 429–442 (2016).

46. Dey, S. et al. ATF4-dependent induction of heme oxygenase 1 prevents anoikis and promotes metastasis. J. Clin. Invest. 125, 2592–608 (2015).

47. Labuschagne, C. F., Cheung, E. C., Blagih, J., Domart, M.-C. & Vousden, K. H. Cell Clustering Promotes a Metabolic Switch that Supports Metastatic Colonization. Cell Metab. 30, 720–734.e5 (2019).

48. Jiang, L. et al. Reductive carboxylation supports redox homeostasis during anchorage-independent growth. Nature 532, 255–8 (2016).

49. Adeshakin, F. O. et al. Mechanisms for Modulating Anoikis Resistance in Cancer and the Relevance of Metabolic Reprogramming. Front. Oncol. 11, 626577 (2021).

50. Gkountela, S. et al. Circulating Tumor Cell Clustering Shapes DNA Methylation to Enable Metastasis Seeding. Cell 176, 98–112.e14 (2019).

51. Rossi, M. et al. PHGDH heterogeneity potentiates cancer cell dissemination and metastasis. Nature 605, 747–753 (2022).

52. Bergers, G. & Fendt, S.-M. The metabolism of cancer cells during metastasis. Nat. Rev. Cancer 21, 162–180 (2021).

53. Mashimo, T. et al. Acetate is a bioenergetic substrate for human glioblastoma and brain metastases. Cell 159, 1603–14 (2014).

54. Bu, P. et al. Aldolase B-Mediated Fructose Metabolism Drives Metabolic Reprogramming of Colon Cancer Liver Metastasis. Cell Metab. 27, 1249–1262.e4 (2018).

55. Christen, S. et al. Breast Cancer-Derived Lung Metastases Show Increased Pyruvate Carboxylase-Dependent Anaplerosis. Cell Rep. 17, 837–848 (2016).

56. Pavlova, N. N. et al. As Extracellular Glutamine Levels Decline, Asparagine Becomes an Essential Amino Acid. Cell Metab. 27, 428–438.e5 (2018).

57. Goodarzi, H. et al. Modulated Expression of Specific tRNAs Drives Gene Expression and Cancer Progression. Cell 165, 1416–1427 (2016).

58. Zhong, Q., et al. Protein posttranslational modifications in health and diseases: Functions, regulatory mechanisms, and therapeutic implications. MedComm (Beijing). 4, e261 (2023).

59. Pavlova, N. N. et al. Translation in amino-acid-poor environments is limited by tRNAGln charging. Elife 9, (2020).

60. Knott, S. R. V et al. Asparagine bioavailability governs metastasis in a model of breast cancer. Nature 554, 378–381 (2018).

61. Rodriguez, E., Lindijer, D. V, van Vliet, S. J., Garcia Vallejo, J. J. & van Kooyk, Y. The transcriptional landscape of glycosylation-related genes in cancer. iScience 27, 109037 (2024).

62. Vuorio, J. et al. N-Glycosylation can selectively block or foster different receptor-ligand binding modes. Sci. Rep. 11, 5239 (2021).

63. Liu, X. et al. Homophilic CD44 Interactions Mediate Tumor Cell Aggregation and Polyclonal Metastasis in Patient-Derived Breast Cancer Models. Cancer Discov. 9, 96–113 (2019).

64. Chanmee, T., Ontong, P., Kimata, K. & Itano, N. Key Roles of Hyaluronan and Its CD44 Receptor in the Stemness and Survival of Cancer Stem Cells. Front. Oncol. 5, 180 (2015).

65. Xu, H., Niu, M., Yuan, X., Wu, K. & Liu, A. CD44 as a tumor biomarker and therapeutic target. Exp. Hematol. Oncol. 9, 36 (2020).

66. Liu, S. et al. CD44 is a potential immunotherapeutic target and affects macrophage infiltration leading to poor prognosis. Sci. Rep. 13, 9657 (2023).

67. Menke-van der Houven van Oordt, C. W., et al. First-in-human phase I clinical trial of RG7356, an anti-CD44 humanized antibody, in patients with advanced, CD44-expressing solid tumors. Oncotarget 7, 80046–80058 (2016).

68. Jiang, J., Batra, S. & Zhang, J. Asparagine: A Metabolite to Be Targeted in Cancers. Metabolites 11, (2021).

69. Juluri, K. R., Siu, C. & Cassaday, R. D. Asparaginase in the Treatment of Acute Lymphoblastic Leukemia in Adults: Current Evidence and Place in Therapy. Blood Lymphat. Cancer 12, 55–79 (2022).

70. Lessner, H. E., Valenstein, S., Kaplan, R., DeSimone, P. & Yunis, A. Phase II study of L-asparaginase in the treatment of pancreatic carcinoma. Cancer Treat. Rep. 64, 1359–61 (1980).

71. Taylor, C. W., Dorr, R. T., Fanta, P., Hersh, E. M. & Salmon, S. E. A phase I and pharmacodynamic evaluation of polyethylene glycol-conjugated L-asparaginase in patients with advanced solid tumors. Cancer Chemother. Pharmacol. 47, 83–8 (2001).

72. Apfel, V. et al. Therapeutic Assessment of Targeting ASNS Combined with l-Asparaginase Treatment in Solid Tumors and Investigation of Resistance Mechanisms. ACS Pharmacol. Transl. Sci. 4, 327–337 (2021).

73. Toth, C. A., Kuklenyik, Z. & Barr, J. R. Nuts and Bolts of Protein Quantification by Online Trypsin Digestion Coupled LC-MS/MS Analysis. Methods Mol. Biol. 1871, 295–311 (2019).

74. Tyanova, S., Temu, T. & Cox, J. The MaxQuant computational platform for mass spectrometry-based shotgun proteomics. Nat. Protoc. 11, 2301–2319 (2016).

75. Tyanova, S. et al. The Perseus computational platform for comprehensive analysis of (prote)omics data. Nat. Methods 13, 731–40 (2016).

76. Chen, S. Ultrafast one-pass FASTQ data preprocessing, quality control, and deduplication using fastp. iMeta 2, e107 (2023).

77. Dobin, A. et al. STAR: ultrafast universal RNA-seq aligner. Bioinformatics 29, 15–21 (2013).

78. Anders, S., Pyl, P. T. & Huber, W. HTSeq--a Python framework to work with high-throughput sequencing data. Bioinformatics 31, 166–9 (2015).

79. Love, M. I., Huber, W. & Anders, S. Moderated estimation of fold change and dispersion for RNA-seq data with DESeq2. Genome Biol. 15, 550 (2014).

80. Yu, G., Wang, L.-G., Yan, G.-R. & He, Q.-Y. DOSE: an R/Bioconductor package for disease ontology semantic and enrichment analysis. Bioinformatics 31, 608–9 (2015).

